# Root metaxylem area influences drought tolerance and transpiration in pearl millet in a soil texture dependent manner

**DOI:** 10.1101/2024.11.09.622826

**Authors:** Pablo Affortit, Awa Faye, Dylan H. Jones, Ezenwoko Benson, Bassirou Sine, James Burridge, Mame Sokhatil Ndoye, Luke Barry, Daniel Moukouanga, Stephanie Barnard, Rahul Bhosale, Tony Pridmore, Pascal Gantet, Vincent Vadez, Philippe Cubry, Ndjido Kane, Malcolm Bennett, Jonathan A. Atkinson, Laurent Laplaze, Darren M. Wells, Alexandre Grondin

## Abstract

- Pearl millet is a key cereal for food security in drylands but its yield is strongly impacted by drought. We investigated how root anatomical traits contribute to mitigating the effects of vegetative drought stress in pearl millet.
- We examined associations between root anatomical traits and agronomical performance in a pearl millet diversity panel under irrigated and vegetative drought stress treatments in field trials. The impact of associated anatomical traits on transpiration was assessed using subpanels grown in different soil within a greenhouse.
- In the field, total metaxylem area was positively correlated with grain weight and its maintenance under drought. In the greenhouse, genotypes with larger metaxylem area grown in sandy soil exhibited a consumerist water use strategy under irrigation, which shifted to a conservative strategy under drought. Water savings was mediated by transpiration restriction under high evaporative demand. This mechanism was dependent on soil hydraulics as it was not observed in peat soil with higher hydraulic conductivity upon soil drying.
- We propose that water savings under drought, mediated by large metaxylem area and its interaction with soil hydraulics, help mitigate vegetative drought stress. Our findings highlight the role of soil hydraulic properties in shaping plant hydraulics and drought tolerance.

## Introduction

Pearl millet is a C4 plant with high nutritional quality that was domesticated about 4,500 years ago in the Sahel region (Burgarella *et al*., 2018). It is the sixth most important cereal crop globally, and its production is concentrated in the arid and semi-arid regions of Western and Central Africa as well as India (Varshney *et al*., 2017). It plays a major role for food security in these regions, serving as a subsistence crop and an important source of revenue for smallholder farmers (FAOSTAT, 2024). Although pearl millet is considered as one of the most heat and drought stress tolerant cereal crops, its production is strongly impacted by climate change (Debieu *et al*., 2017). Pearl millet yield losses due to drought in Sahelian conditions have been estimated at up to 65% during early-stage and vegetative drought stress (i.e. before flowering), with lower losses observed during terminal drought stress (i.e. post flowering; Mahalakshmi *et al*., 1987; Winkel *et al*., 1997). A modelling approach using climatic data collected from 2000 to 2020 in Senegal suggested that vegetative drought stress occurred more frequently than terminal drought stress (24% of the years against 19%, respectively), resulting in a stronger impact on biomass (44% yield loss against 12%, respectively) and grain yield (43% yield loss against 25%, respectively; Fuente *et al*., 2024). In West African traditional farming systems, vegetative drought stress affecting pearl millet often occurs because the crop is typically cultivated under rainfed conditions and sown before or immediately after the first rain, a period characterized by erratic rainfall and potential dry spells lasting several weeks. As current models predict an increase in the frequency of these dry spells in future climate (Sultan & Gaetani, 2016), improving pearl millet’s tolerance to vegetative drought stress is becoming increasingly urgent.

Drought stress occurs when water availability insufficient to meet the plant’s needs for optimal growth. Roots play a critical role in plant water availability and overall plant hydraulics by controlling the access and transport of water to the shoots (Vadez *et al*., 2024). Crops could be bred for root traits that more efficiently and effectively acquire and transport soil water resources, leading to improved drought tolerance (Lynch *et al*., 2021). However, breeding programs have often overlooked root traits in efforts to enhance drought tolerance in crops, because roots are challenging to phenotype in soils and root pre-breeding research is frequently difficult to translate into practical applications in farmer’s fields (Ndoye *et al*., 2022). While technologies for high-throughput phenotyping of roots are advancing (Atkinson *et al*., 2019; Li *et al*., 2022), current research emphasizes field phenotyping of roots (Lynch *et al*., 2021) and aims to better comprehend the complex interplay between root and soil properties on whole plant water uptake (Cai *et al*., 2022). Recent data suggest that as soil dries, stomatal closure is influenced by plant traits related to root hydraulic conductance, as well as by soil texture (Carminati & Javaux, 2020; Cai *et al*., 2022). For instance, maize plants with larger root and rhizosphere systems were able to delay the impacts of soil hydraulic limitations on transpiration in drying soils (Koehler *et al*., 2023), a phenomenon influenced by soil texture (Koehler *et al*., 2022). The ability of pearl millet to thrive in marginal soils with low water retention, while enduring high levels of drought stress, suggests that this species possesses root hydraulic mechanisms that effectively regulate water flow across the soil-plant-atmosphere continuum.

Architectural root traits play a crucial role in determining a plant’s capacity to access water. Deeper root growth is one such trait that helps mitigate the effects of drought stress (Uga *et al*., 2013; Bacher *et al*., 2023). Early pearl millet root growth is characterised by a single primary root being the only architectural component for the first six days (Passot *et al*., 2016). The fast growth of this primary root at depth was associated with improved tolerance to post-germination drought stress under field conditions (Fuente *et al*., 2024). Additional work suggested that, during later stages of pearl millet root development, larger root surface active for water uptake limited the drop in soil water potential around the root and allowed maintenance of transpiration in drying soils (Cai *et al*., 2020). In addition to these architectural traits, root anatomical differences significantly affect plant’s capacity to acquire water under drought stress (Lynch *et al*., 2014; Vadez *et al*., 2024). For example, a reduction in xylem diameter has been shown to improve drought tolerance in wheat by slowing root water uptake and overall plant water use (Richards & Passioura, 1989). Other traits such as root diameter, the presence of cortical aerenchyma, and a reduced number of cortical cell layers, enhance water uptake efficiency by lowering the metabolic cost of root growth (Zhu *et al*., 2010; Chimungu *et al*., 2014a; Sidhu & Lynch, 2024). Moreover, suberization of the endodermis or exodermis layers influences radial water transport (Henry *et al*., 2012; Cantó-Pastor *et al*., 2024). However, little is known about the pearl millet root anatomy, its diversity, and its impacts on root water uptake in relation with soil texture in the context of drought.

In this study, we investigated root anatomical traits in a diverse panel of pearl millet genotypes grown in the field under both irrigated and vegetative drought stress treatments over two consecutive years. We also analysed the variation in shoot biomass and grain weight responses to vegetative drought stress, and linked these to root anatomical traits to determine if any of the later traits were associated with improved drought tolerance. A correlation between total metaxylem area and grain weight and grain weight maintenance under drought was observed. The influence of this trait on transpiration dynamics was further assessed using contrasting subpanels grown in different soil types on a lysimetric platform installed within a greenhouse.

## Material and Methods

### Plant material

One hundred sixty two pearl millet genotypes selected from the pearl millet genetic association panel (PMiGAP; Sehgal *et al*., 2015) were used in this study (Supplementary file **S1**). This panel of inbred lines is composed of cultivated germplasm originating from Africa and India and elite improved open-pollinated cultivars, and is representative of the genetic diversity of pearl millet (Sehgal *et al*., 2015; Varshney *et al*., 2017). We included in this panel Tift23DB that has been used to produce the pearl millet reference genome sequence (Varshney *et al*., 2017; Salson *et al*., 2023), four inbred mapping population parents (ICML-IS11084 and ICML-IS11139 contrasting for soil aggregation, and ICML-IS11155 and SL2 contrasting for primary root length; Fuente *et al*., 2022; Fuente *et al*., 2024) and five inbred lines from West Africa (Debieu *et al*., 2018) and four Senegalese breeding lines. Seeds from the same multiplication were used in both field trials to ensure the genetic uniformity of the genotypes and avoid seed lot effects.

### Field experiments

Pearl millet genotypes were grown in a soil of loamy sand texture (84.8% sand, 7.9% silt and 7.2% clay with a bulk density of 1.7 g cm^-3^ in average from 0 to 200 cm depth; Diongue *et al*., 2022) under irrigated and drought stress treatments over two years (2021 and 2022) in the CNRA research station in Bambey, Senegal (14°42’48.3”N 16°28’41.2”W). The field was laid out using an Alpha Lattice design for each treatment, each including one hundred sixty genotypes with four repetitions or complete blocks that were composed of 10 subblocks. Fifteen genotypes differed from 2021 to 2022 (Supplementary file **S1**). Each subblock included 16 plots and each plot contained 24 hills of one plant of the same genotype (3 rows of eight plants with 90 cm distance between rows and 30 cm distance between plants within the row; Fig. **S1**). Plants were sown in the dry season (early March in 2021 and late March in 2022) to fully control the water supply (weather data are presented in Table **S1**). After sowing, plants were irrigated with 30 mm of water twice a week before application of the drought stress treatment. Drought stress was imposed by withholding water from 21 to 49 days after sowing (DAS) in 2021 and from 21 to 42 DAS in 2022. Both treatments were then irrigated with 30 mm of water twice a week until maturity. Volumetric soil water content was monitored between 0 and 160 cm depth using DIVINER probes (Sentek Pty Ltd) installed throughout the field (Fig. **S2**).

At the end of the drought stress treatment, three representative plants per plot were harvested for crown root anatomical phenotyping following the shovelomics method developed by Trachsel *et al*. (2011) and shoot biomass (SDW) measurement. Crown roots from node four were collected (2 cm segment located approximately 1 cm away from the shoot base) and immediately stored in 50% (v/v) ethanol in water. Roots from this node were chosen because they emerged around 21 DAS, which corresponds to the time of drought stress imposition (Ndoye *et al*., 2024). Furthermore, they were easier to identify and sustained less damage than roots from earlier nodes during sampling.

Phenology, shoot morphology and biomass at maturity, as well as grain weight and yield component traits were measured on three plants per plot as in Debieu *et al*. (2018). The stress tolerance index (STI) defined in Fernandez (1992) was used to evaluate shoot biomass and grain weight (GW) maintenance under drought stress in the panel. This index was calculated on several variables (Var) as :

STI_Var*i*_=(Var_w*i*_* Var_D*i*_)/(Var_w_)²

where Var_w*i*_ is the variable value measured under irrigated condition for genotype *i*, Var_D*i*_ is the variable value measured under drought stress treatment for genotype *i*, Var_w_ is the variable mean value under irrigated condition for all tested genotypes. The STI was measured on grain weight (STI GW), on shoot biomass after the drought stress treatment (STI SDW 49 DAS in 2021 and STI SDW 42 DAS in 2022) and at maturity (STI SDW Mat). STI is a useful metric to compare individual genotypes tolerance to drought while considering its performance under irrigated treatment.

### Crown root anatomical phenotyping

Root anatomical phenotyping was performed using laser ablation tomography (LAT) at the Sutton Bonington campus of the University of Nottingham (UK). Root samples in 50% (v/v) ethanol solution were transferred to custom holders (each holding 48 roots) and moved to a 70% (v/v) ethanol solution for 48 h. Filled holders were then transferred to 100% methanol for a further 48 h before a final transfer to 100% ethanol for a minimum of 48 h prior to critical point drying. Three holders were then transferred to a critical point dryer (Model EM CPD300, Leica Microsystems) and dried by exchange with liquid carbon dioxide. Once dried, holders were stored in containers with silica gel as a desiccant.

LAT uses an ultrafast UV laser to ablate sections from a root sample prior to imaging of the exposed surface, which is illuminated and caused to autofluoresce by the UV laser. Progressing the sample into the ablation plane allows serial imaging of exposed internal surfaces (Chimungu *et al*., 2015; Hall *et al*., 2019; Strock *et al*., 2019; Cunha Neto *et al*., 2023). The tomograph used in this study (LATScan, Lasers for Innovative Solutions LLC) uses a Q-switched UV (355 nm) picosecond-pulsed laser source (Model PX100-3-GF, EdgeWave GmbH) outputting into a galvanometer (Model RTC4, Scanlab GmbH) that focuses the beam via a 160 mm f-theta objective lens. Beam parameters are set by the SPiiPlus software (ACS Motion Control) that also controls a nanopositioning z-stage (PIMag, Physik Instrumente) to advance the sample into the ablation plane. Samples are held in custom holders mounted on XY linear actuators for positioning. Imaging is via a custom system that uses infinity-corrected long working distance objectives (1X – 20X) mounted on a video microscope unit fitted with an objective turret (Model WIDE VMU-H, Mitutoyo (UK) Ltd.) and a 12.3 MP machine vision camera (Model Grasshopper3, FLIR). A third linear actuator, also on the Z plane, positions the imaging system for fine adjustment of focus. An interface written in the LabVIEW development platform (National Instruments Corp.) allows user control of the focus and positioning actuators, camera settings and real-time monitoring of ablation.

An automated sketch in SPiiPlus was used to capture a series of images from critically point dried samples. A series of 10 ablations was performed, first to cut the root at the focal point of the objective, then to ‘polish’ the newly exposed surface by removing any artefacts resulting from thermal ablation, followed by 5 ablations combined with imaging at a z-step size of 10 µm. The sample was then progressed 500 µm into the beam path and the process repeated three times to give a total of 20 images across a 2.6 mm length of root. From the set of 20 imaged sections from each root sample, a representative image was curated for subsequent quantification. These were initially selected from the middle of the image stack, then resampled as necessary to replace images containing features that would hinder analysis, such as points of lateral root emergence, signs of mechanical damage, or uneven laser illumination.

Annotated root images were produced using the CellSeT software tool (Pound *et al*., 2012) to produce a training set for a convolutional neural network for segmentation and quantification based on RootScan and RootScan2 software (Burton *et al*., 2012; https://plantscience.psu.edu/research/labs/roots/methods/computer/rootscan). Nine tissue types consisting of epidermis, sclerenchyma (SCL), cortical cell, aerenchyma, endodermis, stele cell, metaxylem vessels (MX) and vascular bundles were tagged (Fig. **S3**), and an initial set of 200 annotated pearl millet images augmented by rotation, contrast enhancement and degradation, addition of noise, and reflection were used to increase the size of the training set to a total of 1200 images (200 annotated image and 1000 augmented images). The network was created using the PyTorch machine learning framework using a combination of Python and C^++^. Unsupervised training and model generation was run on the University of Nottingham Augusta High Performance Cluster. The training set was supplemented ad hoc by an additional 20 annotated images to improve inaccuracies associated with particular image qualities. The output of the model is a segmented annotated cell mask from which measurements were generated. Features were measured by a combination of pixel and feature counting, and the fitting of circles and convex hulls to the segmentation mask (Fig. **S3b**).

### Crown root anatomical phenotyping along node four in rhizotrons

Root anatomy along the length of crown roots from node four were measured on four genotypes selected for their contrast in total metaxylem area: IP-14210 and IP-5031 are genotypes with small metaxylem area and IP-7536 and IP-6098 are genotypes with large metaxylem area. Seeds were sown in rhizotrons (soil volume -1000 mm x 700 mm x 45 mm) filled with a sandy loam soil (72% sand, 15% silt, 13% clay) irrigated at field capacity in a greenhouse facility at the Sutton Bonington campus of the University of Nottingham (UK) in the 2023 summer season (July-August). Six plants per genotype were planted with one plant per rhizotron for a total of 24 rhizotrons that were randomised in the greenhouse. Temperature was set at 30°C during the day and 24°C at night with 12 hours of artificial lightning from 7AM to 7PM. Plants were well irrigated until harvest at 28 DAS. One whole crown root was then collected on node 4 for each plant and placed in 50% (v/v) ethanol in water. Root samples were collected for LAT imaging at three positions along the root length: at 1 cm from the base of the stem (Base) as in the field experiment, at the middle of the root (Middle) and at 5 cm from the root tip (Apex).

### Transpiration measurements in the greenhouse

Greenhouse experiments were conducted in spring 2023 and 2024 (March-April) at IRD Montpellier (France). In 2023, twelve genotypes contrasted for total metaxylem area but showing no significant differences in terms of shoot biomass measured at 49/42 DAS under irrigated conditions in the field experiments (IP-10539, IP-11929, IP-12395, IP-14210, IP-17150, IP-18168, IP-19626, IP-22494, IP-5031, IP-6098, IP-7536, IP-9496) were cultivated under irrigated and drought stress treatments in an organic potting substrate (referred hereafter as peat soil) in 5.5-liter pots (19 cm top diameter, 15.5 cm bottom diameter and 25 cm depth). The potting substrate (GO M2 140 substrate, Jiffy Group) was composed of crushed baltic white peat (75%), coconut peat (20%) and sand (5%). It was implemented with optimal fertilisation for pearl millet growth. Pots were placed on load cells (240 in total) installed within the greenhouse. The load cells were set to monitor pot weight every 30 min (Phenospex Ltd). The temperature in the greenhouses was set at 28°C during the day and 25°C at night. Daylength was controlled by artificial lighting (900 W m^-2^) from 7AM to 7PM. Relative humidity, temperature and solar radiation were monitored in the greenhouse. Genotypes were completely randomised with six replicates per treatment. In each replication, both treatments were positioned side by side in the greenhouse to limit positioning effects. In 2024, twenty genotypes contrasting for the same criteria (IP-11929, IP-12395, IP-14210, IP-17150, IP-18168, IP-19626, IP-22494, IP-5031, IP-6098, IP-7536, IP-9496 from the 2023 experiment implemented with IP-10759, IP-10964, IP-12364, IP-15533, IP-17611, IP-21206, IP-22420, IP-22423, IP-5272) were grown in similar conditions except that the soil used was a mix at equivalent volume (50:50 v/v) of the previous potting substrate with fine sand (AF0/1RS, DIALL) of particle size below 1 mm (referred hereafter as sandy soil).

Plants were well irrigated until 20 DAS. At this date, pots were saturated and drained overnight. At 21 DAS, a layer of plastic beads was added on the soil surface in order to prevent soil evaporation and pots weight were measured to record the weight at field capacity. An experiment performed beforehand following a similar set up was conducted for each soil to determine the coefficient of pot weight reduction from field capacity to soil water content at which the plant stopped transpiring. These coefficients (70% and 30% reduction of the initial pot weight for the peat soil and for the sandy soil, respectively) were used to approximate, from the pots saturated weights, the pots weight at which plant transpiration theoretically stopped. The difference between the weight at field capacity and the weight at which transpiration stopped corresponded to the fraction of transpirable soil water (FTSW). FTSW was further used to monitor soil water content through daily recordings of pots weight at 3PM. The drought stress treatment of the peat soil experiment consisted in a dry down to FTSW 40% (pot weight when 40% of the water usable for transpiration is left) that was maintained till harvest (42 DAS). The drought stress treatment of the sandy soil experiment consisted in a progressive dry-down so all plants reached FTSW 75% (at 28 DAS), then FTSW 60% ( at 35 DAS) and FTSW 40% (at 40 DAS), after which irrigation was stopped until harvest (42 DAS). In the irrigated treatment of both experiments, soil moisture was maintained at FTSW 75% until harvest.

At harvest, leaves were collected to measure leaf area using a planimeter (LI-3100C, LI-COR). Leaves and stems were further oven dried for three days at 70°C to measure dry biomass. For each plant, crown roots from nodes one to five in the peat soil experiment and from nodes three and four in the sandy soil experiment were collected and stored in 2 ml tubes filled with 50% (v/v) ethanol in water for LAT. Root theoretical axial hydraulic conductance (*K*_x_ in m^4^ s^−1^ MPa^−1^) was estimated as the sum of the theoretical axial root hydraulic conductance of each individual metaxylem vessels present in the root cross section using the following modified Hagen-Poiseuille equation:

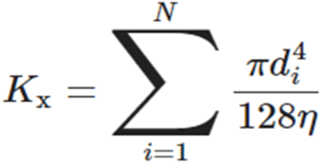

where *N* is the number of xylem vessels in a root cross section, *d* is the diameter of one individual xylem vessel in the root cross section and *η* is the viscosity of water (1.0021 × 10^−9^ MPa s^-1^ at 20°C; Nobel, 2009).

Slopes of the transpiration response to the evaporative demand were calculated at 41 DAS in both greenhouse experiments using an automated pipeline described by Kar *et al*. (2020). Briefly, plant transpiration (Tr) was first calculated as the difference in pot weights within intervals of 30 minutes. Transpiration rate (Tr Rate) was further measured by dividing Tr by the time interval and plant leaf area, assuming marginal changes in leaf area between 41 and 42 DAS. At each time interval, reference evapotranspiration (ETref) was calculated according to the Penman-Monteith equation using actual climatic data within the greenhouse (Zotarelli *et al*., 2010). Slopes of transpiration response to the evaporative demand (Slope Tr) were calculated from 7AM to 3:30PM (which corresponded to the maximum evaporative demand of the day) by plotting the Tr Rate against the ETref. Profiles of Tr along the day were plotted for two groups of genotypes contrasting for their theoretical axial root hydraulic conductance (*K*_x_).

### Data analysis

Outliers within each genotype were statistically removed using the interquartile method in R for all measured traits in each experiment (ggplot2 v0.6.0 boxplot. stats et grDevices v4.3.3). Data obtained from each field experiment were further corrected for spatial heterogeneity by treatments using the Spatial Analysis of Trials using splines (SpATs) function in the StatgenSTA package in R (Rodríguez-Álvarez *et al*., 2018; van Rossum, 2024), considering a resolvable incomplete block design model. In this model, genotype was fitted as a fixed effect and Best Linear Unbiased Estimates (BLUEs) were produced. Heritability, considered as the ratio of genetic variance to phenotypic variance, was calculated in StatgenSTA for each trait using a similar model but fitting the genotype as a random effect. To correct for spatial heterogeneity in the greenhouse experiments, a similar approach was followed but using the SpATS model in the StatgenHTP package in R (Millet *et al*., 2021) which allows spatial correction on individual plants.

### Statistical analysis

Statistical analyses were carried out in R (V 4.3.3; R Development Core Team, 2008). Normality of the different traits were verified using the Shapiro-Wilk test and the equality of variances using the Levene test. Trait means between treatments within years were compared using the Wilcoxon test. The Student t-test was used to compare mean values of total metaxylem area in the rhizotron experiment and transpiration (Tr) in the greenhouse experiments. Correlations between variables were investigated using the Pearson correlation test using BLUEs.

## Results

### Vegetative drought stress negatively impacts shoot biomass production and grain weight in pearl millet

Field trials were conducted over for two consecutive years (2021 and 2022) at CNRA Bambey in Senegal, using diverse pearl millet genotypes. Genotypes were cultivated either under irrigated treatment or subjected to vegetative drought stress by withholding irrigation 21 days after sowing. The drought stress lasted four weeks (from 21 to 49 DAS) in 2021 and three weeks (from 21 and 42 DAS) in 2022. After the stress period, crown roots from node four were sampled for anatomical phenotyping and shoot biomass was measured. Irrigation was then resumed until maturity, at which agro-morphological and yield component traits were measured to study the diversity of drought responses within the panel.

A large variability in all agronomic traits was observed, under both irrigated and drought stress treatments (Table **1**). Heritability ranged from 0.4 for shoot biomass measured at 49 DAS under drought in 2021 to 0.9 for days to flowering in both years and treatments, indicating that these traits are under strong genetic control (Table **1**). Furthermore, positive and significant correlations were observed for both agro-morphological and yield component traits between the two years under irrigated and drought treatments (Fig. **S4**), illustrating the robustness of the measured traits from one year to the next.

**Table 1.** Variation for morphological and agronomical traits in the field trials.

Mean: Mean of n = 4. SD: Standard deviation; CV: Coefficient of variation; $H^2$ : Heritability.
| Acronym | Trait | Year | Treatment | Mean | SD | CV | $H^2$ |
| --- | --- | --- | --- | --- | --- | --- | --- |
| GW | Grain weight (g plant<sup>-1</sup>) | 2021 | WW | 89 | 39 | 1488 | 0.5 |
|  |  |  | WS | 43 | 26 | 694 | 0.6 |
|  |  | 2022 | WW | 71 | 36 | 1262 | 0.7 |
|  |  |  | WS | 46 | 28 | 810 | 0.6 |
| DTF | Days to 50% flowering | 2021 | WW | 58 | 7.5 | 56 | 0.9 |
|  |  |  | WS | 65 | 9.2 | 83 | 0.9 |
|  |  | 2022 | WW | 64 | 11 | 115 | 0.9 |
|  |  |  | WS | 71 | 13 | 182 | 0.9 |
| SDW | Shoot biomass at the end of the stress (g plant<sup>-1</sup>) | 2021 | WW | 55 | 28 | 769 | 0.6 |
|  |  |  | WS | 20 | 12 | 154 | 0.4 |
|  |  | 2022 | WW | 27 | 15 | 221 | 0.7 |
|  |  |  | WS | 10 | 5.8 | 34 | 0.6 |
| SDW Mat | Shoot biomass at maturity (g plant<sup>-1</sup>) | 2021 | WW | 169 | 71 | 5017 | 0.7 |
|  |  |  | WS | 87 | 52 | 2717 | 0.7 |
|  |  | 2022 | WW | 196 | 93 | 8675 | 0.8 |
|  |  |  | WS | 151 | 78 | 6080 | 0.8 |
| 1000-grain weight | 1000 grains weight (g plant<sup>-1</sup>) | 2021 | WW | 8.5 | 1.6 | 2.66 | 0.6 |
|  |  |  | WS | 7.4 | 1.7 | 2.8 | 0.8 |
|  |  | 2022 | WW | 8.6 | 2.2 | 4.8 | 0.8 |
|  |  |  | WS | 8.3 | 1.9 | 3.57 | 0.8 |
| Tiller number | Number of tillers | 2021 | WW | 9.8 | 4 | 15.65 | 0.5 |
|  |  |  | WS | 8.3 | 3.9 | 15.19 | 0.7 |
|  |  | 2022 | WW | 9.9 | 3.4 | 11.3 | 0.7 |
|  |  |  | WS | 8.7 | 2.7 | 7.41 | 0.7 |
| Plant height | Plant height at maturity (cm) | 2021 | WW | 155 | 29 | 856 | 0.9 |
|  |  |  | WS | 122 | 31 | 956 | 0.8 |
|  |  | 2022 | WW | 144 | 28 | 776 | 0.9 |
|  |  |  | WS | 131 | 34 | 1131 | 0.9 |

Drought stress significantly reduced shoot biomass measured at 49 and 42 DAS in both years, with a 60% and 61% average reduction in 2021 and 2022, respectively (Fig. **1a**). At maturity, partial recovery in shoot biomass was observed with average reduction of 46% in 2021 and 25% in 2022 (Fig. **1b**). However, grain weight was reduced by 53% in 2021 and 37% in 2022 (Fig. **1c**). The drought stress also had a negative and significant impact on plant height, the number of tillers, and thousand grain weight in both years (Fig. **S5**).

**Fig. 1.**
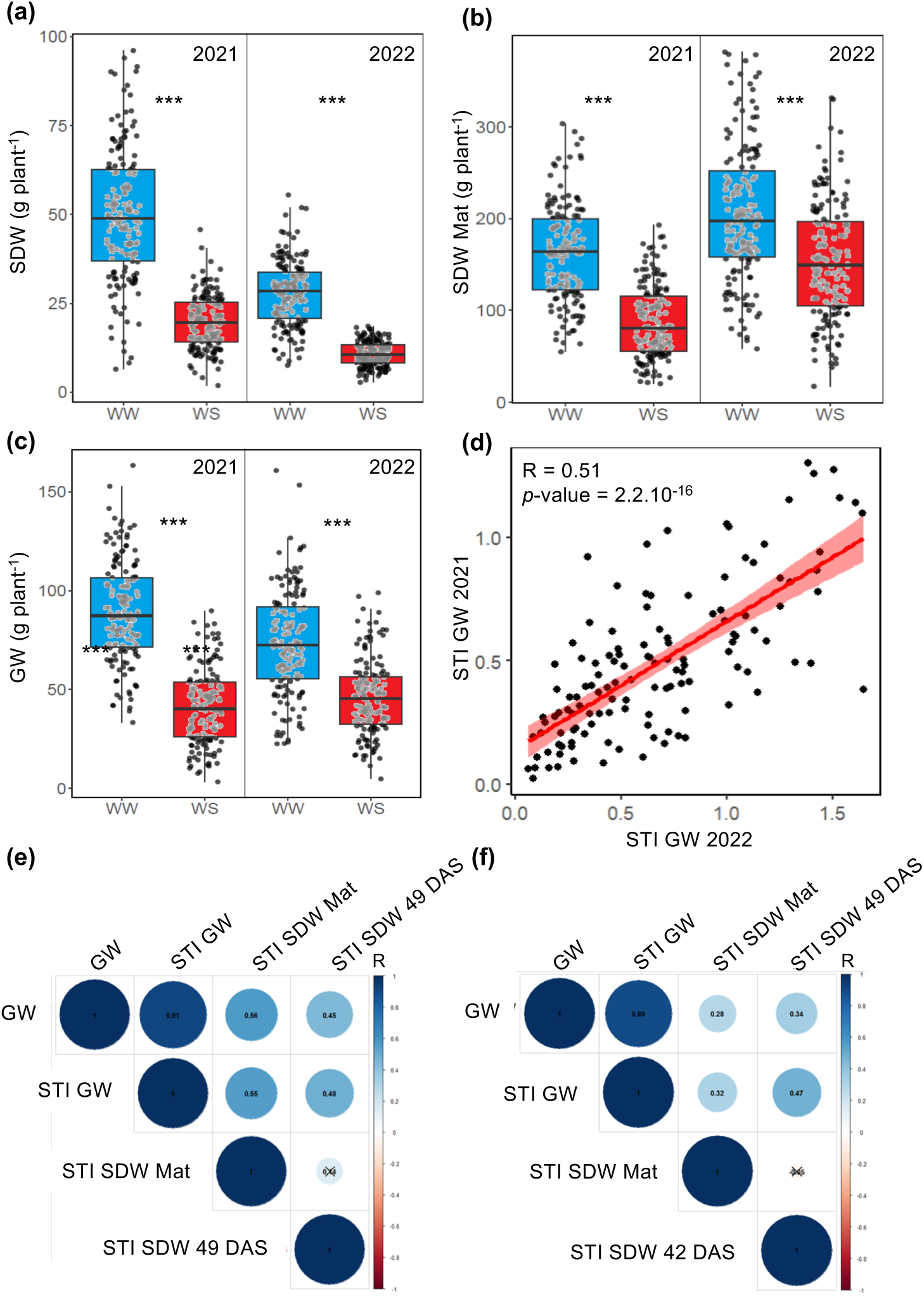
Drought stress effects on shoot biomass and grain yield in the field. (a) Shoot biomass at the end of the drought stress (SDW 49 DAS in 2021 and SDW 42 DAS in 2022), (b) shoot biomass at maturity (SDW Mat), and (c) grain yield (GW) were measured in the field on diverse pearl millet genotypes under well-watered (WW) and drought stress (WS) treatments in 2021 and 2022. *** *p-*value < 0.001 according to a Wilcoxon test. (d) Covariation between stress tolerance index for grain yield (STI GW) measured in 2021 and 2022. The Pearson correlation coefficient (R) and *p*-value of the correlation test are indicated. (e, f) Correlation between grain weight (GW), the stress tolerance index for shoot biomass measured at the end of the drought stress (STI SDW 49 DAS in 2021 and STI SDW 42 DAS in 2022), the shoot biomass measured at maturity (STI SDW Mat) and the stress tolerance index for grain yield (STI GW) in 2021 (e) and 2022 (f). Boxplots and correlation analyses were performed using the best linear unbiased estimates. The Pearson correlation coefficient (R) is indicated for each pair of traits and black crosses indicate non-significant correlation (*p-*value > 0.05).

The stress tolerance index (STI) was used to measure maintenance of shoot biomass at the end of the drought stress period, as well as the maintenance of shoot biomass and grain weight at maturity. This index corresponds to an indicator of drought tolerance and performance under irrigated conditions, both of which are crucial for breeding purposes. Large variation in stress tolerance index for both shoot biomass measured at the end of the drought stress and at maturity, and for grain weight, was observed in each year, with values significantly correlated between years (R = 0.51 for grain weight; Fig. **1d** and **S6**). The stress tolerance index for grain weight varied from low values (0.02 at lowest in 2021 and 2022) for genotypes performing poorly under both irrigated and drought stress treatments to values above 1 (1.3 at highest in 2021 and 1.8 in 2022) for genotypes performing well under both treatments. Furthermore, the stress tolerance index for grain weight was positively correlated with the stress tolerance index for shoot biomass measured at the end of the drought stress and at maturity in both years (Fig. **1e**,**f**), suggesting that maintaining shoot biomass production under stress was associated with grain weight maintenance.

Altogether, the field trials revealed a large phenotypic diversity for shoot morphological and agronomical traits in pearl millet. Vegetative drought stress had a significant impact on most traits, with genotypes better able to maintain shoot biomass production during the stress period showing improved grain weight maintenance under drought.

### Metaxylem area positively associates with grain weight production and maintenance under early drought stress

To identify root anatomical traits that contribute to grain weight and its maintenance under vegetative drought stress in pearl millet, cross-sectional images of crown roots sampled from node four at the end of the drought stress in the field were obtained using laser ablation tomography (LAT) and anatomical traits were measured using an updated version of RootScan for pearl millet (Fig. **2a,b****,c**). Traits of the epidermis, cortical cells, aerenchyma, endodermis and vascular bundle were excluded due to lower measurement reliability.

**Fig. 2.**
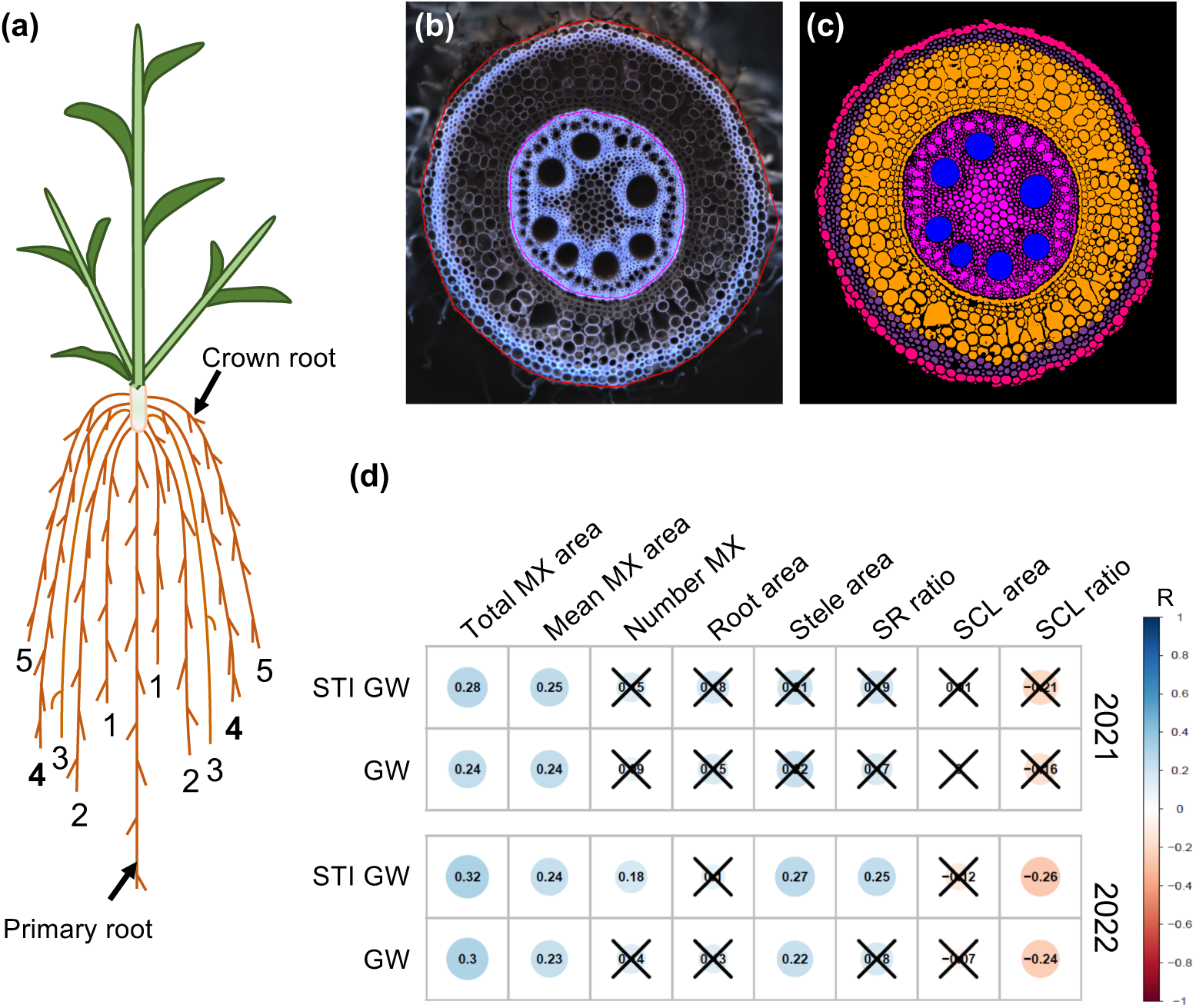
Correlation between root anatomical traits and yield. (a) Schematic representation of a pearl millet root system with crown roots from different nodes. (b) Image of a pearl millet root cross section obtained through laser ablation tomography. Root cross section area (Root area; red line) and stele area (Stele area; pink line) were estimated using the convex hull approach. (c) Same image as in (b) segmented using RootScan adapted for pearl millet. Metaxylem (MX) were segmented in blue pixel and resulted in the quantification of total metaxylem area (sum of the blue pixels), number of metaxylem (number of blue objects) and mean area of one metaxylem vessel (total metaxylem area divided by the number of metaxylem). Stele was segmented in pink. Cortex was segmented in orange. Sclerenchyma (SCL) area was quantified as the sum of the purple pixels. Epidermis was segmented in reddish purple. (d) Correlation between grain weight (GW) measured under drought stress, the stress tolerance index for grain weight (STI GW) and root anatomical traits measured under drought stress in the 2021 and 2022 field experiments. Correlation test was performed using the best linear unbiased estimates. The Pearson correlation coefficient (R) is indicated for each pair of traits and black crosses indicate non-significant correlation (*p*-value > 0.05). SCL ratio: ratio of sclerenchyma area to root cross section area; SR ratio: ratio of stele area to root cross section area.

Large variation in for total metaxylem area, number of metaxylem vessels, mean area of individual metaxylem vessels, root cross section area, stele area, sclerenchyma area, ratio between stele area and root cross section area, and the ratio between sclerenchyma area and root cross section area were observed within the panel under both treatments (Table **2**). The heritability of these traits varied from 0.4 for total metaxylem area measured in the 2021 drought stress treatment to 0.81 for mean area of individual metaxylem vessels measured in the 2022 irrigated treatment (Table **2**). Positive and significant correlations were observed for root anatomical traits between years under irrigated treatment (Fig. **S7**). Under drought stress, a similar trend was observed, but, correlations between years were weaker and only significant for root cross section area, stele area and metaxylem-related traits (Fig. **S7**). Within years, drought stress significantly negatively impacted all measured root anatomical traits in 2021, but only affected stele area and the ratio between stele area and root cross section area in 2022 (Fig. **S8**).

**Table 2.**
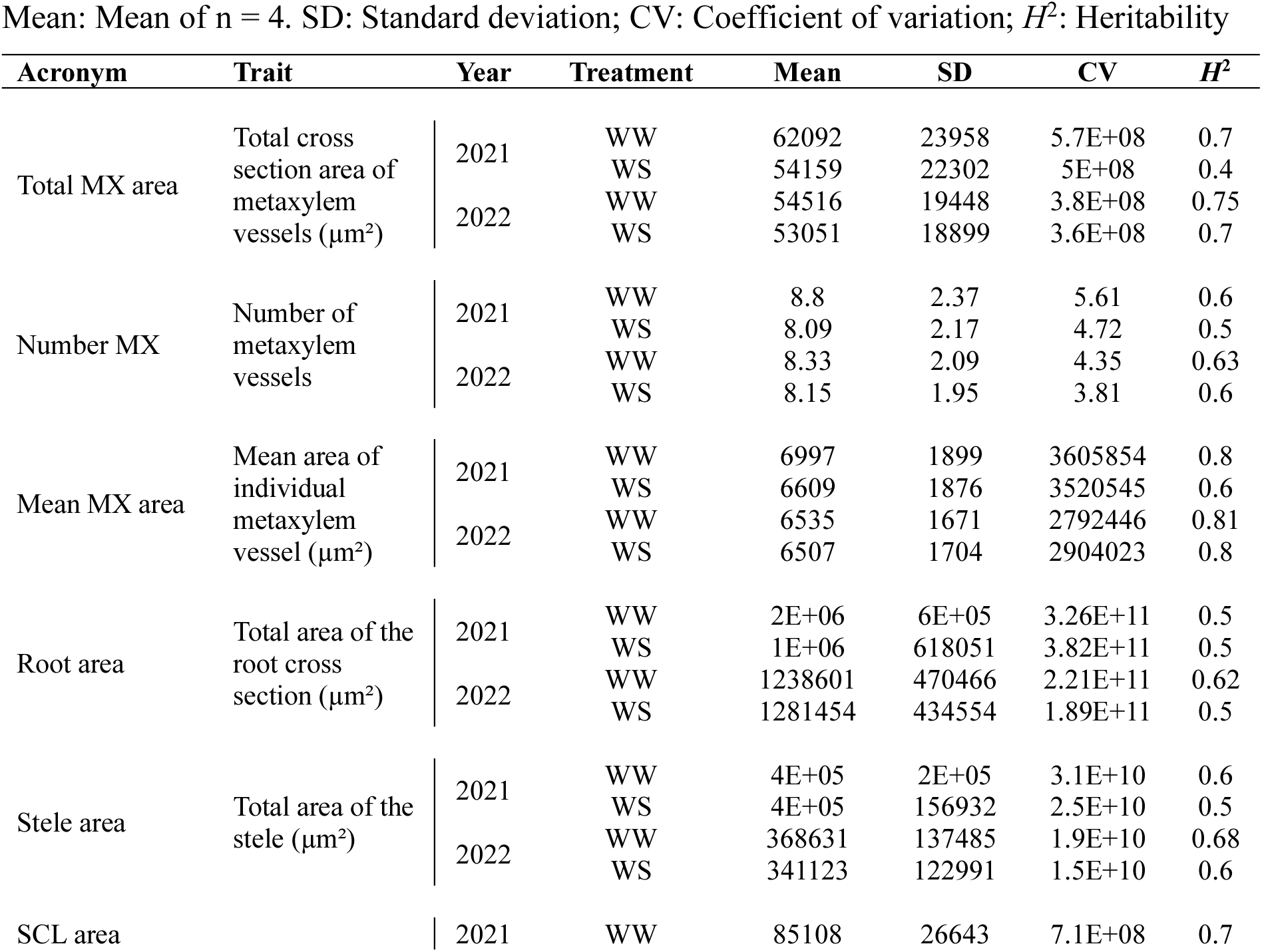
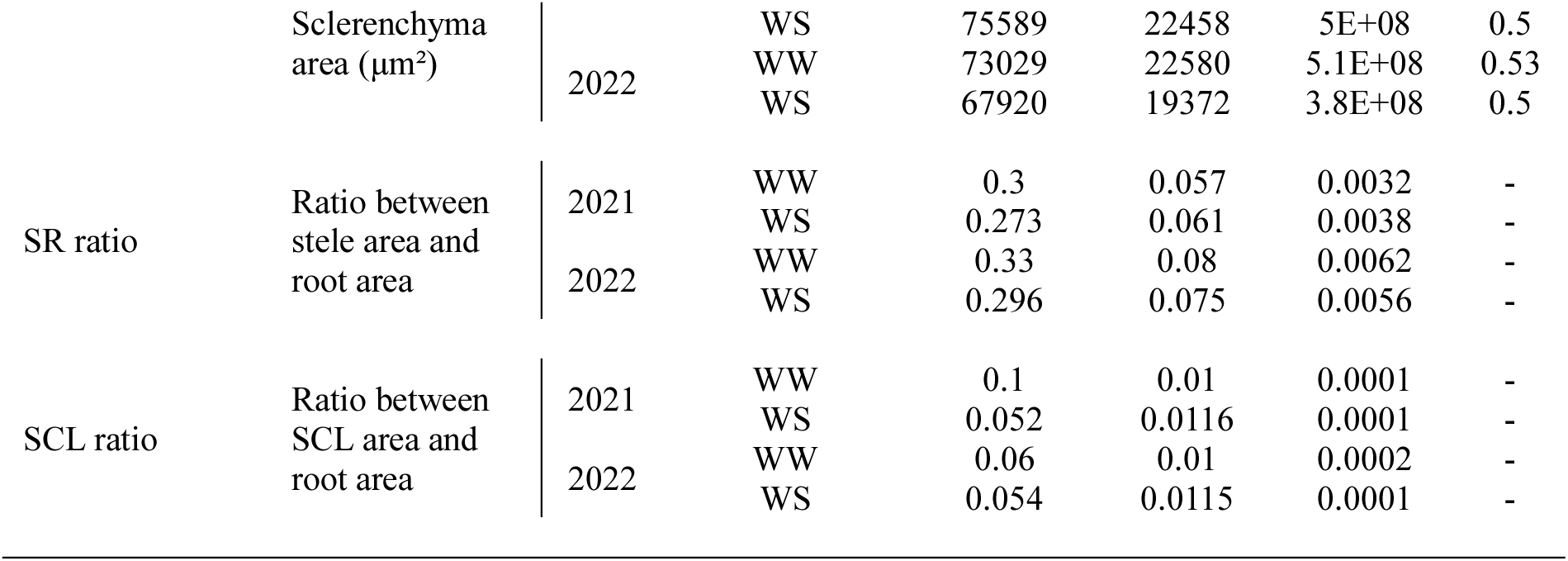
Variation for root anatomical traits in the field trials.

We further investigated correlations between root anatomical traits, shoot biomass, grain weight and the stress tolerance indices. Positive and significant correlations were observed in both years between total metaxylem area and grain weight measured under drought stress, and stress tolerance index for grain weight (Fig. **2d**). Similar correlations were observed between mean area of individual metaxylem vessels and grain weight measured under drought stress, and stress tolerance index for grain weight (Fig. **2d**). These metaxylem traits measured under irrigated treatment were also correlated with grain weight measured in the same condition and the stress tolerance index for grain weight in both years (Fig. **S9**). This suggests that pearl millet genotypes with larger total metaxylem area were both more productive under irrigated treatment and more tolerant in terms of grain weight under vegetative drought stress. Positive and significant correlations were also observed between the stele area and the ratio between stele area and root cross section area measured under drought stress and the stress tolerance index for grain weight, although only significant in 2022 (Fig. **2d**). Similarly, negative and significant correlations between the ratio of sclerenchyma area and root cross section area measured under drought stress and stress tolerance index for grain weight were observed only in 2022 (Fig. **2d**).

Hence, several root anatomical traits were associated with grain weight under both irrigated and drought stress treatments in pearl millet. Among them, total metaxylem area and mean area of metaxylem vessels were the most consistently positively associated with grain weight under both treatments and its maintenance under drought.

### Metaxylem area measured on crown roots from node four are representative of the overall plant’ root metaxylem characteristics

In the field trials, root anatomy was studied on crown roots from node four sampled near the stem base. To determine whether metaxylem characteristics were conserved along the crown root from node four, we measured root anatomy at three different locations in four genotypes contrasting for total metaxylem area grown in rhizotron (Fig. **3a**). The contrast for total metaxylem area observed in the field between genotypes at the stem base was conserved, with IP-14210 having the smaller total metaxylem area and IP-6098 having the larger total metaxylem area (Fig. **3b**). Total metaxylem area measured at the base of the stem where metaxylem vessels are fully elongated, was not significantly different from that measured at the middle of the root, except in IP-14210 (Fig. **3b**). However, total metaxylem area at these two locations was significantly greater than that measured at the root apex, with the exception of IP-14210. In fact, no contrast between genotypes was observed for total metaxylem area at the root apex. The number of metaxylem vessels also correlated strongly from the stem base to the root apex (Fig. **S10a**). These results suggest that variations between metaxylem-related traits between pearl millet genotypes appear in the elongated root region and that variations in these traits are likely similar along the root.

**Fig. 3.**
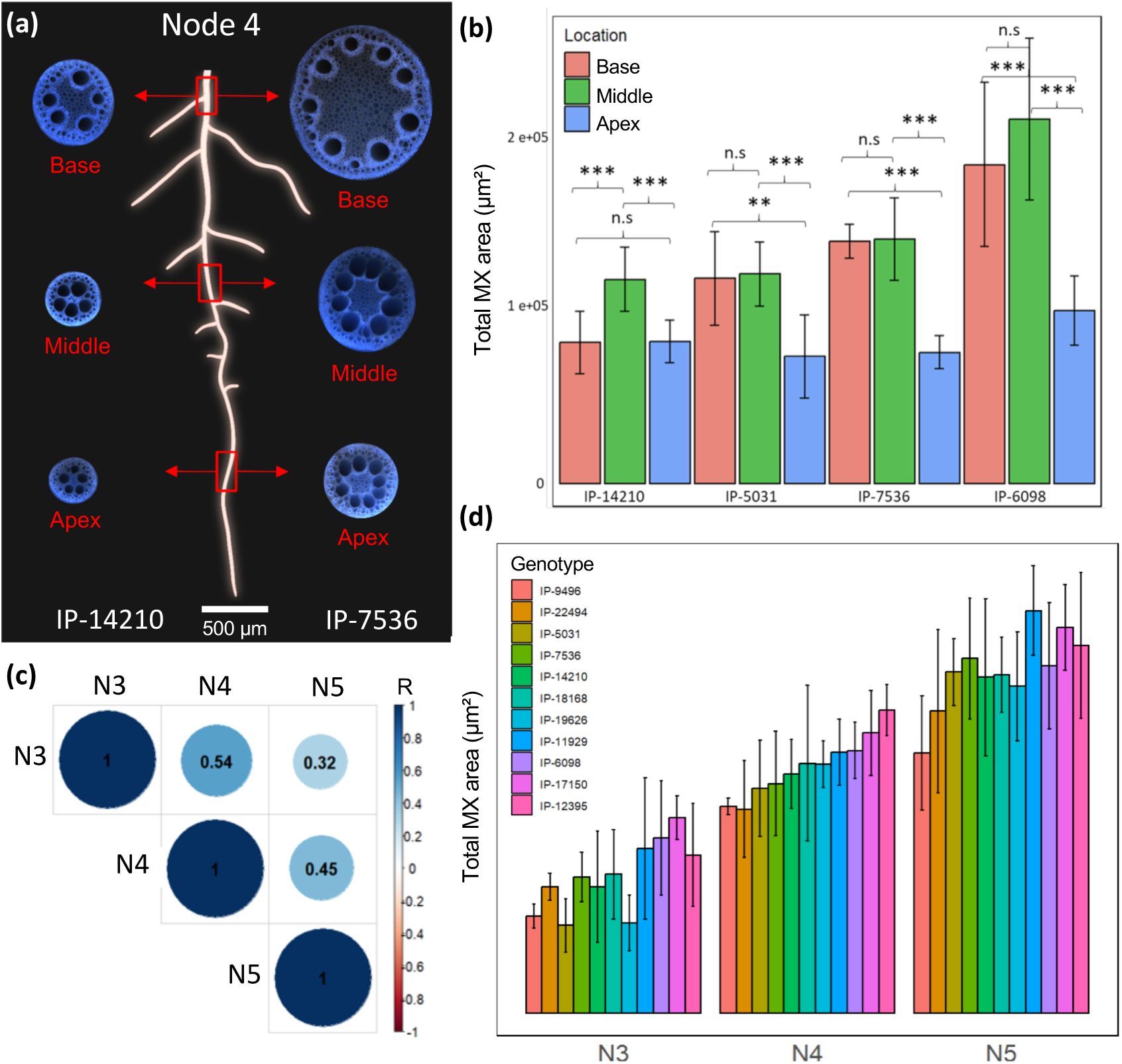
Total metaxylem area along the crown root from node 4 and in different nodes. (a) Images of root anatomy at three locations (base, middle and apex), along the root length of a crown root sampled from node four in two contrasted genotypes for total metaxylem (MX) area. (b) Total metaxylem area measured at the base, middle and apex of crown roots from node four in four contrasted genotypes for this trait. Plants were grown in rhizotrons under irrigated conditions. ** *p*-value < 0.01, *** *p*-value < 0.001 according to a Student t-test. n.s.: Non-significant. Bars represent means of n = 6 root cross-sectional images per location ± se. (c) Correlation between the total metaxylem area measured at the base of crown roots from node three, four and five. Plants were grown in peat soil under irrigated treatment in the 2023 greenhouse experiment. The Pearson correlation coefficient (R) is indicated for each pair of traits. (d) Genotypes ranking for total metaxylem area measured at the base of crown roots from node three(N3), four (N4) and five (N5). Genotypes were ranked based on their total metaxylem area measured on node four. Plants were grown in the same conditions as in (c). Bars represent means of n = 6 root cross-sectional images per node ± se.

We next tested whether total metaxylem area measured on crown root from node four was representative of total metaxylem area from roots of earlier and later nodes in 12 genotypes, which differed in their total metaxylem area. These genotypes were grown in pots filled with peat soil in a greenhouse. We aimed to assess total metaxylem area in crown roots from nodes one and two, but, these roots were very thin, and anatomical measurements using LAT were challenging. Total metaxylem area was therefore measured in roots from nodes three, four, and five. Total metaxylem area and the number of metaxylem vessels from all nodes were positively and significantly correlated (Fig. **3c** and Fig. **S10b**), except for the number between node three and node five. The strongest correlations were observed between total metaxylem area of crown roots from adjacent nodes (nodes four and five), while the weakest correlations were observed between total metaxylem area of crown roots from nodes further apart (nodes three and five; Fig. **3c**). Total metaxylem area of crown roots increased from node three to five, but the ranking of genotypes was generally conserved from one node to another, indicating the variation in total area of metaxylem of crown roots is proportional across nodes within genotypes (Fig. **3d**).

Altogether, these results suggest that total metaxylem area measured in crown roots at the base of the stem on node four is representative of the overall root metaxylem characteristics of the plant. A plant with a smaller total metaxylem area in crown roots at the stem base of node four will likely show a smaller total metaxylem area along the crown root in the fully elongated zone and across crown roots from different nodes, and vice versa.

### Axial root hydraulic conductance correlates with transpiration restriction under high evaporative demand in dry sandy soil

A positive correlation between total metaxylem area, grain weight and the maintenance of grain weight under vegetative drought stress was observed in the field experiments. As metaxylem vessel size determines axial root conductance, we hypothesised that total metaxylem area could impact transpiration and plant water use under drought stress. To test if total metaxylem area influences transpiration, we monitored the transpiration of genotypes with contrasting total metaxylem areas, grown under irrigated and gradual vegetative drought stress treatments in pots filled with sandy soil in a greenhouse, as evaporative demand increases. At 41 DAS, plant transpiration (Tr) was measured along the day and plant transpiration rate (TR Rate) was plotted against the ETref in order to measure the transpiration response to the evaporative demand, as the slope of the linear regression between these two variables (Slope TR; Fig. **S11**).

The total metaxylem area varied significantly between genotypes (Fig. **S12a**) and significant effect of the drought stress was observed on shoot biomass, indicating that the drought stress was effectively sensed by the plants (Fig. **S12b**). Drought stress did not significantly affect the total metaxylem area and the mean area of individual metaxylem vessels (Fig. **S12c,d**), indicating no drought-induced plastic responses of these traits in this subpanel. Theoretical axial root hydraulic conductance (*K*_x_) was strongly correlated with total metaxylem area (Fig. **S13**) and further used in correlation analyses with plant transpiration as it represents a more physiologically relevant variable. Under irrigated treatment, no correlations were observed between the axial root hydraulic conductance of crown roots from nodes three and four, and the slope of transpiration response to the evaporative demand (Fig. **S14a,b**). Conversely, in the drought stress treatment a negative and significant correlation between the axial root hydraulic conductance of crown roots from node four and the slope of transpiration to the evaporative demand was observed (Fig. **4b**). A similar negative trend, although not significant, was observed with the axial root hydraulic conductance of crown roots from node three (Fig. **4a**). This suggests that under drought stress, plants with larger axial root hydraulic conductance grown in sandy soil restricted their transpiration more when the evaporative demand was the highest, compared to plants with smaller axial root hydraulic conductance.

**Fig. 4.**
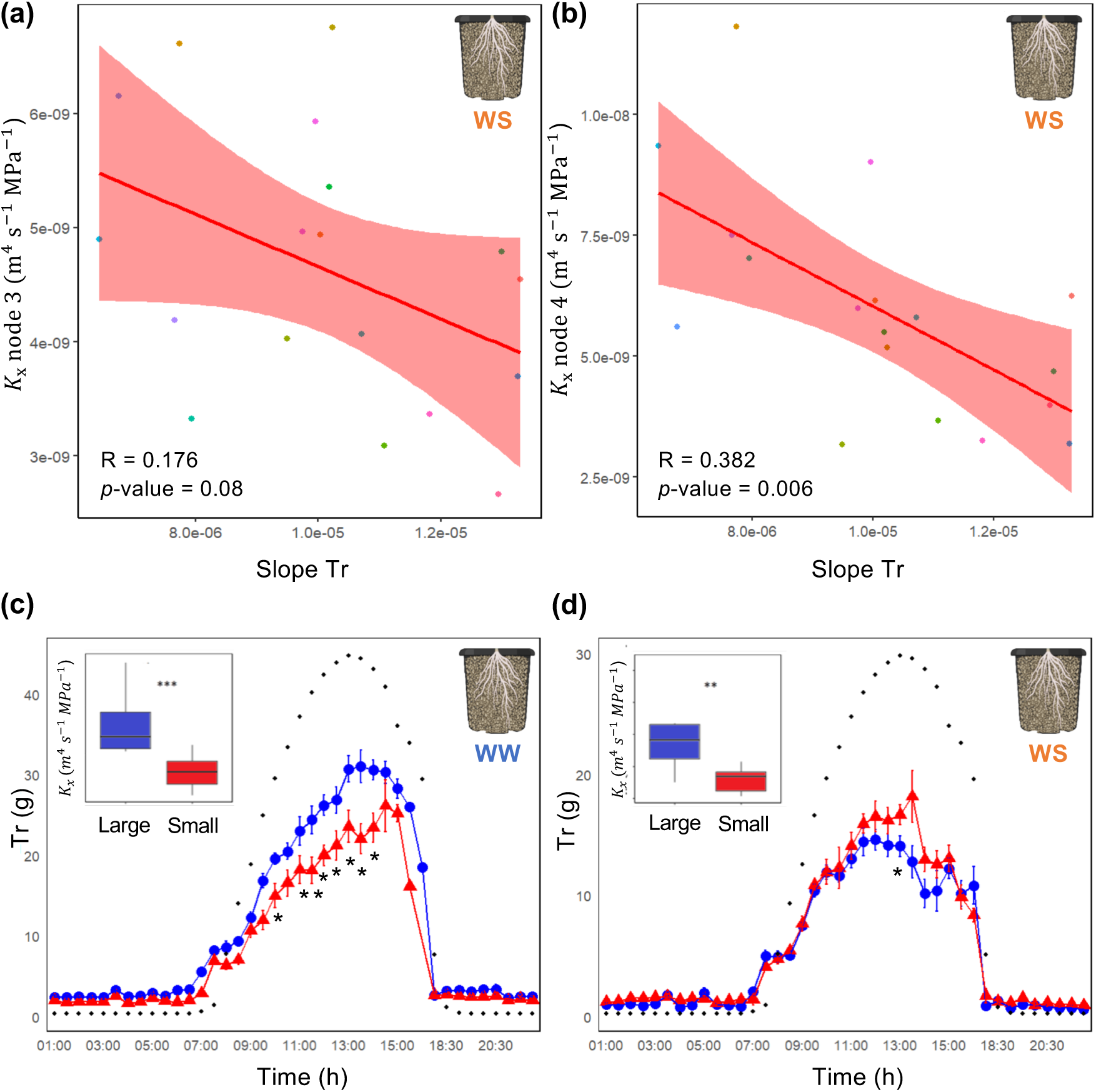
Association between axial root hydraulic conductance and transpiration response to the evaporative demand in sandy soil. (a, b) Covariation between axial root hydraulic conductance (*K*_x_)measured on crown roots from node three (a) and four (b), and the slope of the transpiration response to the evaporative demand, both measured under drought stress (WS) treatment in the greenhouse. Pearson correlation coefficient (R) and *p*-value of the correlation test are indicated. (c, d) Transpiration (Tr) along the day in two groups of genotypes contrasted for axial root hydraulic conductance (large in blue and small in red)measured under irrigated (c) and drought stress treatments (d). Black dots represent the change in reference evapotranspiration (ETRef) along the day. Significant differences in Tr between the two groups were assessed using a Student t-test. * *p*-value < 0.05.

Higher axial root hydraulic conductance translated into more water use when the water was not limited, as illustrated by the significant positive and significant correlation between the cumulated water use at 41 DAS and the axial root hydraulic conductance of crown roots from node four in the irrigated treatment (Fig. **S15a**). Conversely, when the water was limited, genotypes with higher axial root hydraulic conductance used significantly less water than genotypes with lower axial root hydraulic conductance, as shown by the significant negative correlation between the cumulated water use at 41 DAS and the axial root hydraulic conductance of crown roots from node four in drought stress (Fig. **S15b**). Daily transpiration profiles for two groups of genotypes contrasting for axial root hydraulic conductance but not for shoot biomass (Fig. **S16a**) showed that these responses occur particularly during the hottest hours of the day (Fig. **4c,d**).

Hence, we conclude that, in sandy soil, plants with larger total metaxylem area and higher axial root hydraulic conductance restrict their transpiration during the hottest hours of the day, thus using less water than plants with smaller total metaxylem area and lower axial root hydraulic conductance. We hypothesise that this mechanism leads to water savings under drought stress.

### Soil texture influences the impact of axial root hydraulic conductance on plant transpiration under drought

While our results suggest that larger total metaxylem area are associated with water savings in pearl millet grown under drought in sandy soil, opposite results have been observed in other species (Richards & Passioura, 1989; Salih *et al*., 1999; Hendel *et al*., 2021). Recent studies suggest that soil hydraulic properties influence plant transpiration upon soil drying, as well as plant hydraulic traits (Koehler *et al*., 2023). We therefore tested if the link between total metaxylem area and transpiration restriction at high evaporative demand under drought was conserved in peat soil with different hydraulic properties than sandy soil (higher water retention and hydraulic conductivity as the soil dries; Cai *et al*., 2022; Wankmüller *et al*., 2024), using a similar experimental set up in the greenhouse.

In peat soil, the total metaxylem area also varied significantly between genotypes (Fig. **S17a**) and significant effect of the drought stress was observed on shoot biomass but not on the total metaxylem area and the mean area of individual metaxylem vessels (Fig. **S17b, c, d**). A significant positive correlation between the slope of the transpiration response to the evaporative demand and the axial root hydraulic conductance of crown roots from node three was observed in the irrigated treatment, while a similar but non-significant trend was found for crown roots from node four (Fig. **S14c**,**d**). Under drought stress, a significant positive correlation between the slope of transpiration response to the evaporative demand and the axial root hydraulic conductance of crown roots from node three was observed (Fig. **5a**). Although not significant, the same positive trend was observed on crown roots from node four (Fig. **5b**). Transpiration was significantly higher at the hottest hours of the day for genotypes with higher axial root hydraulic conductance in both treatments (Fig. **5c,d**). However, no significant correlations were observed between the cumulated water use and axial root hydraulic conductance when considering all the genotypes in both treatments (Fig **S18a,b**). No significant differences in shoot biomass were observed between these two groups (Fig. **S16b**). This indicates that in the peat soil, plants with larger axial root conductance were more responsive to the evaporative demand under both irrigated and drought stress treatments.

**Fig. 5.**
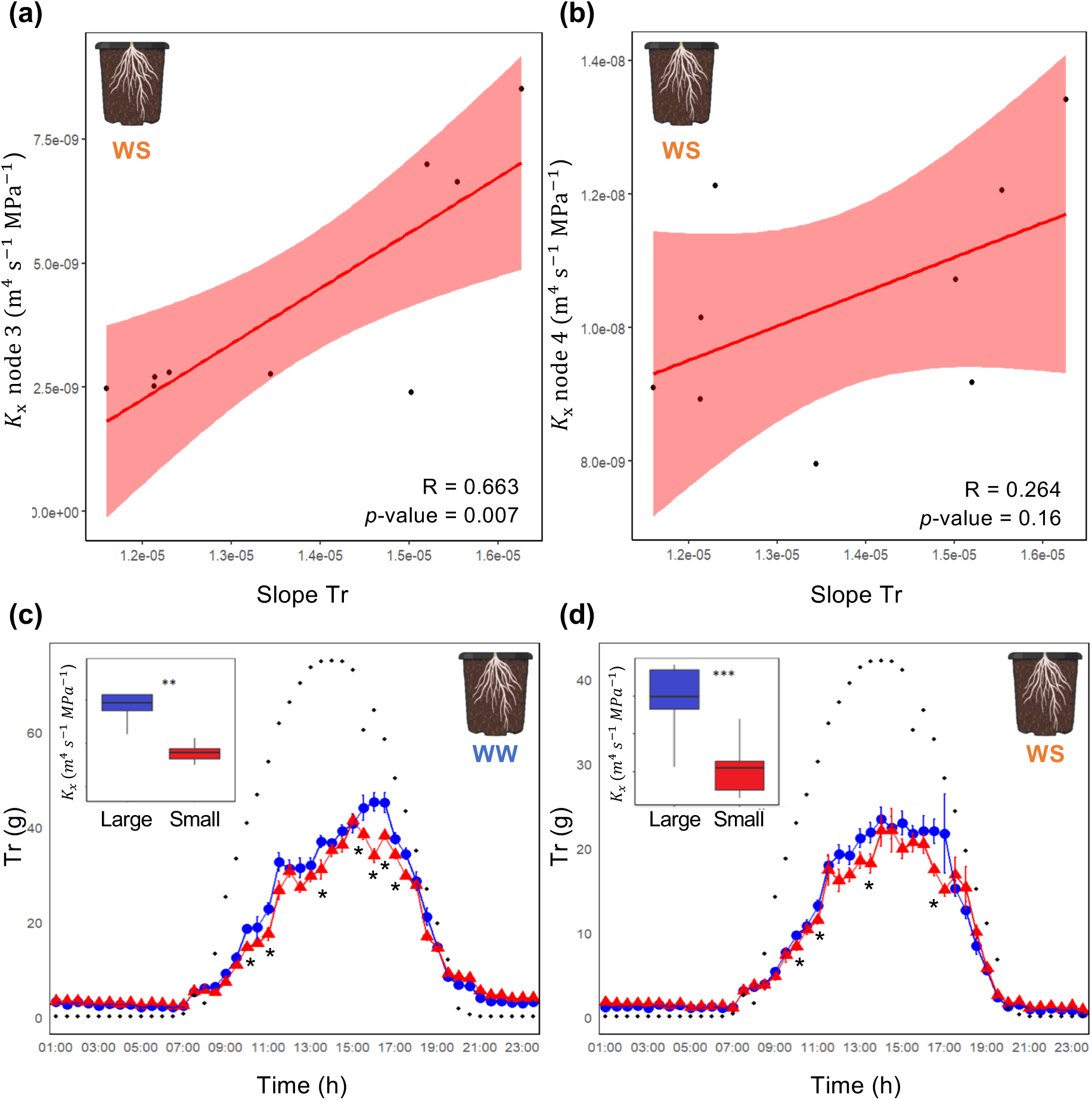
Association between axial root hydraulic conductance and transpiration response to the evaporative demand in peat soil. (a, b) Covariation between axial root hydraulic conductance (*K*_x_) measured on crown roots from node three (a) and four (b), and the slope of the transpiration response to the evaporative demand, both measured under drought stress (WS) treatment in the greenhouse. Pearson correlation coefficient (R) and *p*-value of the correlation test are indicated. (c, d) Transpiration (Tr) along the day in two groups of genotypes contrasted for axial root hydraulic conductance (large in blue and small in red) measured under irrigated (c) and drought stress treatments (d). Black dots represent the change in reference evapotranspiration (ETRef) along the day. Significant differences in Tr between the two groups were assessed using a Student t-test. * *p*-value < 0.05.

Overall, our results suggest that total metaxylem area and axial root hydraulic conductance influence plant transpiration under drought in a soil-dependent manner. Plants with larger total metaxylem area and axial root hydraulic conductance were able to extract more water from peat soil than from sandy soil under drought, and the opposite was true for plants with lower total metaxylem area.

## Discussion

In this study, we report that traits related to total metaxylem area and mean area of metaxylem were consistently associated with grain weight and its maintenance in pearl millet under vegetative drought stress in the field. Measurements of transpiration dynamics in a greenhouse on a subpanel of genotypes contrasting for total metaxylem area showed that genotypes with larger total metaxylem area on crown roots from nodes three and four (representative of total metaxylem area in roots from subsequent nodes), tended to use more water than genotypes with smaller total metaxylem area under irrigated treatment. Under drought, genotypes with larger total metaxylem area restricted transpiration at the hottest hours of the day and used less water than genotypes with smaller metaxylem vessels in a sandy soil. The transpiration response to evaporative demand observed in sandy soil was reversed in peat soil, which has higher hydraulic conductivity than sandy soil during dry down conditions.

In maize, anatomical phenotyping of roots on node four proved effective in investigating the significance of traits such as cortical cell size and root cortical aerenchyma in drought tolerance (Chimungu *et al*., 2014b, 2015) or multiseriate cortical aerenchyma in soil compaction (Schneider *et al*., 2021). In pearl millet, we also phenotyped roots on node four as its development was initiated around the time of drought stress imposition (Ndoye *et al*., 2024). Our results suggest that total metaxylem area was conserved along the root from the elongation zone to the oldest root base tissue. In line with previous results obtained in maize (Yang *et al*., 2019), total metaxylem area correlated positively between nodes three to five in eleven pearl millet genotypes contrasting for total metaxylem area, suggesting that anatomical observations of metaxylem-related traits on node four are representative of metaxylem features in subsequent nodes.

Water uptake mainly occurs through crown roots and their laterals in maize (Ahmed *et al*., 2016, 2018). Similar processes likely occur in pearl millet that does not display seminal roots (Passot *et al*., 2016) and in which the primary root degenerates a few weeks after germination (ICRISAT, 1981). Water uptake is determined by the speed at which water can be channelled from the soil to the root, i.e. the root hydraulic conductance which can be decomposed into radial and axial conductance. Although radial conductance may represent the most obvious limitation to root hydraulic conductivity, a compelling body of evidence suggest that axial conductance can also be limiting to water uptake in certain plants and environmental conditions (Passioura, 1983; Doussan *et al*., 1998; Couvreur *et al*., 2012; Javaux *et al*., 2013). A limitation comes from the diameter of the vessels which was formalised by Hagen-Poiseuille’s law, vessels with lower diameter having lower hydraulic conductance. In maize primary roots, positive correlation was observed between the number and area of metaxylem vessels and the root hydraulic conductivity (Rishmawi *et al*., 2023). Furthermore, modelling showed that root system conductance was sensitive to axial transport in wheat (Bouda *et al*., 2018). Considering that metaxylem characteristics were conserved across crown roots, we hypothesised that genotypes with larger metaxylem area on crown root from node four displayed enhanced root hydraulic conductance, with subsequent effects on root water uptake effects and drought tolerance. Although correlations observed between total metaxylem area and drought tolerance measured as grain weight maintenance were significant and robust in the field experiments, the coefficient of correlation was relatively low which may reflect the complexity of different root architectural and anatomical trait interactions on root hydraulics (Maurel & Nacry, 2020). Further architectural phenotyping of genotypes contrasting for total metaxylem area identified in this study should allow better characterization of root hydraulic architecture in pearl millet.

Plant hydraulics may be modelled as a demand and supply network. Stomatal closure and decreased transpiration will occur when the water supply from the roots becomes insufficient to sustain the demand by the shoot (Vadez *et al*., 2024). In wet soils, plants with lower root hydraulic conductance would therefore be more rapidly limited in their ability to supply water to the shoots when the evaporative demand increases, which would lead to reduction in transpiration and subsequent water savings (Passioura, 1983). A breeding program in wheat focussed on reducing xylem diameter and the associated axial root hydraulic conductance resulted in deceleration of water use and improved yield in rain-fed environments prone to terminal drought (Richards & Passioura, 1989). Our study also highlights a link between total metaxylem area and water use in pearl millet. Under irrigated treatment, larger total metaxylem area was associated with higher water use as expected. However, larger total metaxylem area was associated with transpiration restriction and decreased water use when the evaporative demand increased under drought stress in sandy soil. To explain this result, we extended the plant supply and demand hydraulic model mentioned above to the soil hydraulics which can have strong impacts on transpiration under drought (Carminati & Javaux, 2020). In maize, comparison of transpiration response to drought stress using hydraulically contrasting soils showed that transpiration decreased at more negative water potential in loam than in sand (Koehler *et al*., 2022). This response was linked to an abrupt decrease in soil-root hydraulic conductance at less negative water potential in sand (Koehler *et al*., 2022). Indeed, sandy soil displays a lower water retention and a steeper decrease in hydraulic conductivity due to their large pores and narrow pore size distribution compared to loam (Cai *et al*., 2022). It was suggested that while high root hydraulic conductance enables plants to meet high transpiration demand in irrigated soils, it also increases their sensitivity to soil drying in a soil dependent manner (Cai *et al*., 2022; Wankmüller *et al*., 2024) - i.e. plants with high root conductance would sense better a decline in soil hydraulic conductivity. Therefore, the relationship between transpiration response to evaporative demand and total metaxylem area that we observed under drought in sandy soil could be linked to the higher sensitivity to decreasing soil conductivity of the more conductive plants.

We propose that, in sandy soils, larger axial root hydraulic conductance due to larger metaxylem area causes a relatively larger drop in leaf water potential relative to the average leaf water potential, leading to a reduction in soil-root conductance triggering stomatal closure and transpiration restriction. We hypothesise that water refilling of the root-soil interface will occur during the night from wet soil not in contact with roots or in contact with roots that are not active. Therefore, the transpiration restriction response would repeat and potentially amplify in subsequent days of soil drying till the fraction of transpirable soil water is null. As the hydraulic properties of the sandy soil used in this study is representative of soil where pearl millet is grown in West Africa, we suggest that similar mechanisms have occurred during the field trials.

Transpiration restriction in response to increased evaporative demand is a mechanism that allows plants to save water at the hottest hours of the day when water loss is poorly rewarded by carbon fixation (Vadez *et al*., 2023). For this reason, constitutive expression of this phenotype has been linked to increased plant transpiration efficiency (Affortit *et al*., 2022) or tolerance to terminal drought stress with limited trade-offs (Vadez *et al*., 2013; Cooper *et al*., 2014; Sinclair *et al*., 2017). We propose that the transpiration restriction and subsequent water savings observed in genotypes with larger total metaxylem area under drought in sandy soil contributed to mitigate the impacts of the stress. By saving water, these genotypes would have been better able to maintain their shoot biomass during the drought stress period through improved plant transpiration efficiency. Furthermore, these genotypes were also those better able to maintain their shoot biomass and grain weight at harvest. Yet, other mechanisms such as plant vigour or flowering time may have contributed to the improved drought tolerance within the panel. A clustering analysis into genotypes with low and high shoot biomass at the end of the drought stress shows that the association between total metaxylem area and grain weight maintenance under drought stress is stronger in the higher shoot biomass cluster which also shows earlier flowering time (Fig. **S19** and **S4**). This suggests that larger total metaxylem area would be particularly beneficial for vegetative drought stress tolerance in vigorous and early flowering genotypes. However, shoot biomass and grain weight measured in the irrigated treatment correlated poorly with these same traits measured in the drought stress treatment (Fig. **S20**), suggesting that vigour was not the main factor explaining drought tolerance. Furthermore, drought stress tended to delay flowering within the panel suggesting no significant drought escapism effects, and no correlation between flowering time and grain weight and its maintenance was observed (Fig. **S4**). These observations support our conclusions that total metaxylem area significantly influenced vegetative drought stress tolerance in pearl millet.

In West Africa, pearl millet landrace variability has already been exploited by farmers to shorten flowering cycles of existing landraces as a response of the 1970s and 1980s drought episodes (Vigouroux *et al*., 2011; Dussert *et al*., 2015). Souna3, a popular variety in this region that has been selected for drought tolerance, was included in our field trials. This variety showed average total metaxylem area when compared to the phenotypic diversity observed in the panel, indicating that this trait could still be improved in pearl millet. However, this phenotype may not prove beneficial for all types of drought stress. Plants with larger total metaxylem area will use more water under wet conditions which may enhance the risk of water limitations in case of a terminal drought stress. Further characterization of the genetic determinants controlling metaxylem size should produce genetic materials useful to test these hypotheses in field conditions.

## Conclusion

Pearl millet is a cereal crop adapted to arid and semi-arid regions where rain patterns are often erratic and vegetative drought stress can greatly affect grain weight. In sandy soil where pearl millet is typically grown in West Africa, we observed that larger total metaxylem area has benefits for grain weight under irrigated and vegetative drought stress treatments. We propose that larger metaxylem allows increased transpiration when the water is sufficient and water savings through transpiration restriction when water is limited. The hydraulic properties of the sandy soil in which pearl millet is grown may, in relation with plant hydraulics influenced by metaxylem vessel size, be responsible for this shift in water use strategies. Our work reveals the opportunistic nature of pearl millet in terms of water use and highlights how soil hydraulic properties interact with plant hydraulics to influence transpiration along the soil-plant-atmosphere continuum.

## Acknowledgements

We are grateful to Ghislain Kanfany from ISRA and to Prakash Gangashetty and Mohammed Riyazaddin from ICRISAT for providing seeds of the PMiGAP. We thank the members of the CERAAS and CNRA centres of ISRA for their help during the field experiments in Senegal, and members of the CERES team of the UMR DIADE for their support in the greenhouse experiments in Montpellier. We acknowledge Miranda Niemec for helpful advice on root phenotyping, and Jonathan Lynch, Andrea Carminati and Mathieu Javaux for helpful discussions in the preparation of this manuscript.

## Funding

This work was supported by the Royal Society (Anatomics grant ICA-R1-180356 to MB and NK), the USAID Feed the Future Sorghum and Millet Innovation lab (GenMil grant n°S19182.01 to NK), the French Institute for Sustainable Development (IRD), the French Ministry for Research and Higher Education (PhD grant to PA) and the Agence National pour la Recherche (PlastiMil grant ANR-17-CE20-0022 to AG).

## Competing interest

The Authors declare no competing interest.

## Author contributions

Design of the research: PA, RB, TP, PG, VV, PC, NK, MB, DW, LL, JA, AG; Performance of the research: PA, AF, DJ, EB, BS, JB, MSN, LB, DM, MB, SBK, DW, LL, JA, AG; Data analysis, collection or interpretations: PA, AF, DJ, EB, BS, JB, LB, VV, PC, NK, MB, DW, LL, JA, AG; Writing the manuscript: PA, DJ, MB, DW, LL, JA, AG.

## Data availability

All datasets are available upon request.

## Supporting Information

**Table S1** Weather data collected during the field trials in 2021 and 2022.

**Supplementary file S1** Passport data of the different pearl millet lines used in this study.

**Fig. S1** Field trials experimental design.

**Fig. S2** Volumetric soil water content in the 2021 and 2022 field trials at two depth intervals (0-60 cm and 60-120 cm) measured using DIVINER probes.

**Fig. S3** An example cross-sectional image of a crown root from node four obtained through laser ablation tomography and tissue annotation using the pearl millet version of RootScan.

**Fig. S4** Correlation between shoot morphological and agronomical traits measured in the field across both years (2021 and 2022) in the irrigated (WW) and drought stress (WS) treatments.

**Fig. S5** Stress impact on plant height, tiller number and 1000-grain weight measured in the field across both treatments (WW: Irrigated; WS: Drought stress) and years (2021 and 2022).

**Fig. S6** Covariation between stress tolerance index for shoot biomass measured in both years (2021 and 2022).

**Fig. S7** Correlation between root anatomical traits measured in the field across both years (2021 and 2022) in the irrigated (WW) and drought stress (WS) treatments.

**Fig. S8** Stress impact on total metaxylem vessel area, mean area of metaxylem, number of metaxylem, root cross section area, stele area, ratio of stele area to root cross section area (SR ratio), sclerenchyma area and ratio of sclerenchyma to root cross section area (SCL ratio).

**Fig. S9** Correlation between root anatomical traits, grain weight (GW), and stress tolerance index for grain weight (STI GW) measured under irrigated treatment in the 2021 and 2022 field experiments.

**Fig. S10** Correlation between the number of metaxylem vessels along the crown root of node four and across different nodes.

**Fig. S11** Transpiration response to the evaporative demand in pearl millet genotypes grown in the greenhouse under irrigated (WW) and drought stress (WS) treatments.

**Fig. S12** Stress impact on shoot biomass and metaxylem-related traits in pearl millet genotypes contrasting for total metaxylem area.

**Fig. S13** Covariation between total area of metaxylem and axial root hydraulic conductance.

**Fig. S14** Covariation between axial root hydraulic conductance (*K*x) measured in crown roots from node three and four, and the transpiration response to the evaporative demand (Slope Tr) in pearl millet genotypes contrasting for total metaxylem area grown under irrigated treatment in the greenhouse.

**Fig. S15** Water use in pearl millet genotypes contrasting for total metaxylem area grown under irrigated (WW) and drought stress (WS) treatments in sandy soil in the greenhouse.

**Fig. S16** Shoot biomass in two groups of genotypes contrasting for axial root hydraulic conductance (*K*x; large versus small) measured under the drought stress treatment.

**Fig. S17** Stress impact on shoot biomass and metaxylem-related traits in pearl millet genotypes contrasting for total metaxylem area. Plants were grown in the greenhouse in peat soil.

**Fig. S18** Water use in pearl millet genotypes contrasting for total metaxylem area grown under irrigated (WW) and drought stress (WS) treatments in peat soil in the greenhouse.

**Fig. S19** Covariation between total metaxylem area and stress tolerance index for grain weight.

**Fig. S20** Covariation of shoot biomass and grain yield between both treatments within years.

## Notes

### Competing Interest Statement

The authors have declared no competing interest.

