## Supplemental Table and Figures for "Root metaxylem area influences drought tolerance and transpiration in pearl millet in a soil texture dependent manner"

**Table S1** Weather data collected during the field trials in 2021 and 2022. Tmin : Minimum temperature; Tmax : Maximum temperature; HRmin: Minimum relative humidity; Hrmax: Maximum relative humidity; CCI : Cloud cloudiness index; Wind speed at 2 m height from the soil. Data indicate the average over the indicated month. Wind speed was not measured in April and May 2021, and March 2022 due to an instrument failure.

| Year | Month | Rainfall (mm) | Tmin (°C) | Tmax (°C) | Hrmin (%) | Hrmax (%) | CCI (hour) | Wind (m/s at 2m) | Evapotranspiration (mm) |
|---|---|---|---|---|---|---|---|---|---|
| 2021 | March | 0 | 18±0.3 | 37.8±0.4 | 26.9±1.5 | 75.5±2.5 | 9.1±0.3 | 3.2±0.3 | 9.9±0.4 |
| | April | 0 | 19.9±0.2 | 39.8±0.5 | 25.4±1.3 | 75.9±2.4 | 10.2±0.2 | | 11.0±0.5 |
| | May | 0 | 20.4±0.3 | 37.6±0.5 | 31.7±1.6 | 84.6±1.7 | 9.2±0.2 | | 7.8±0.4 |
| | June | 37.3±1.2 | 23.8±0.3 | 37.3±0.4 | 48.0±1.6 | 89.2±1.2 | 7.3±0.6 | 2.1±0.1 | 6.2±0.3 |
| | July | 23.2±1.7 | 24.5±0.2 | 36.4±0.4 | 58.2±1.2 | 91.6±0.7 | 7.7±0.5 | 2.4±0.1 | 5.1±0.2 |
| 2022 | March | 0 | 18.2±0.4 | 36.7±0.4 | 32.8±1.4 | 80.3±2.6 | 9.4±0.2 | | 9.5±0.5 |
| | April | 0 | 20.4±0.4 | 38.1±0.6 | 34.6±1.4 | 81.0±2.8 | 10.1±0.1 | 3.0±0.2 | 9.9±0.7 |
| | May | 2.3±0.1 | 22.1±0.3 | 39.2±0.6 | 34.7±2.4 | 84.1±2.1 | 9.2±0.3 | 2.1±0.1 | 8.4±0.5 |
| | June | 40.1±0.8 | 24.9±0.2 | 36.8±0.5 | 46.3±1.9 | 88.0±1.4 | 6.9±0.6 | 2.4±0.2 | 6.2±0.4 |
| | July | 131.9±1.7 | 25.5±0.2 | 34.4±0.5 | 57.3±2.2 | 90.5±0.9 | 6.8±0.6 | 0.6±0.0 | 4.2±0.3 |

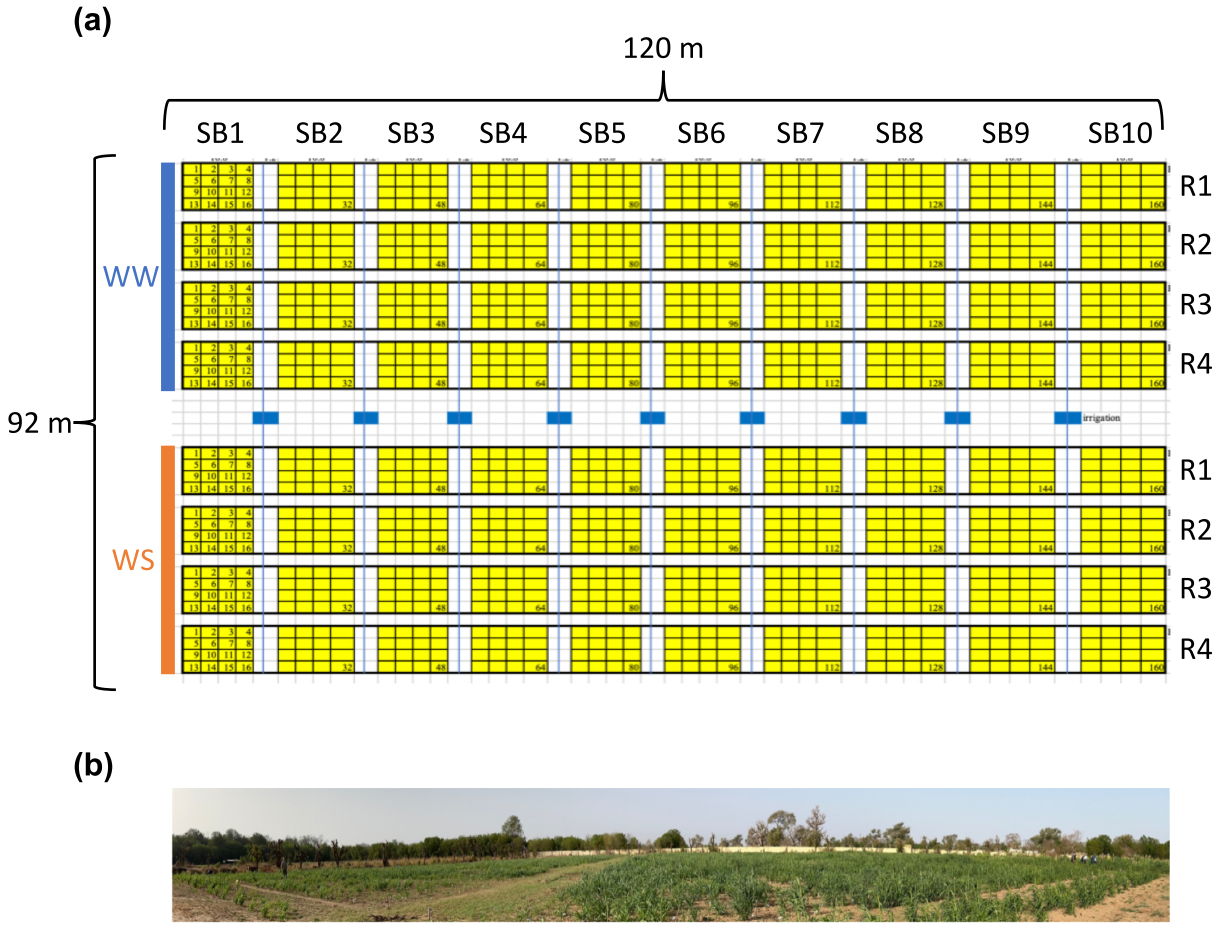


**Fig. S1** Field trials experimental design. (a) Layout with the two treatments (WW: Irrigated; WS: Drought stress), each being composed of four replicates (R) or complete blocks (160 genotypes in total per repetitions). Each block was composed of ten subbocks (SB). (b) Image of the 2022 field trial.

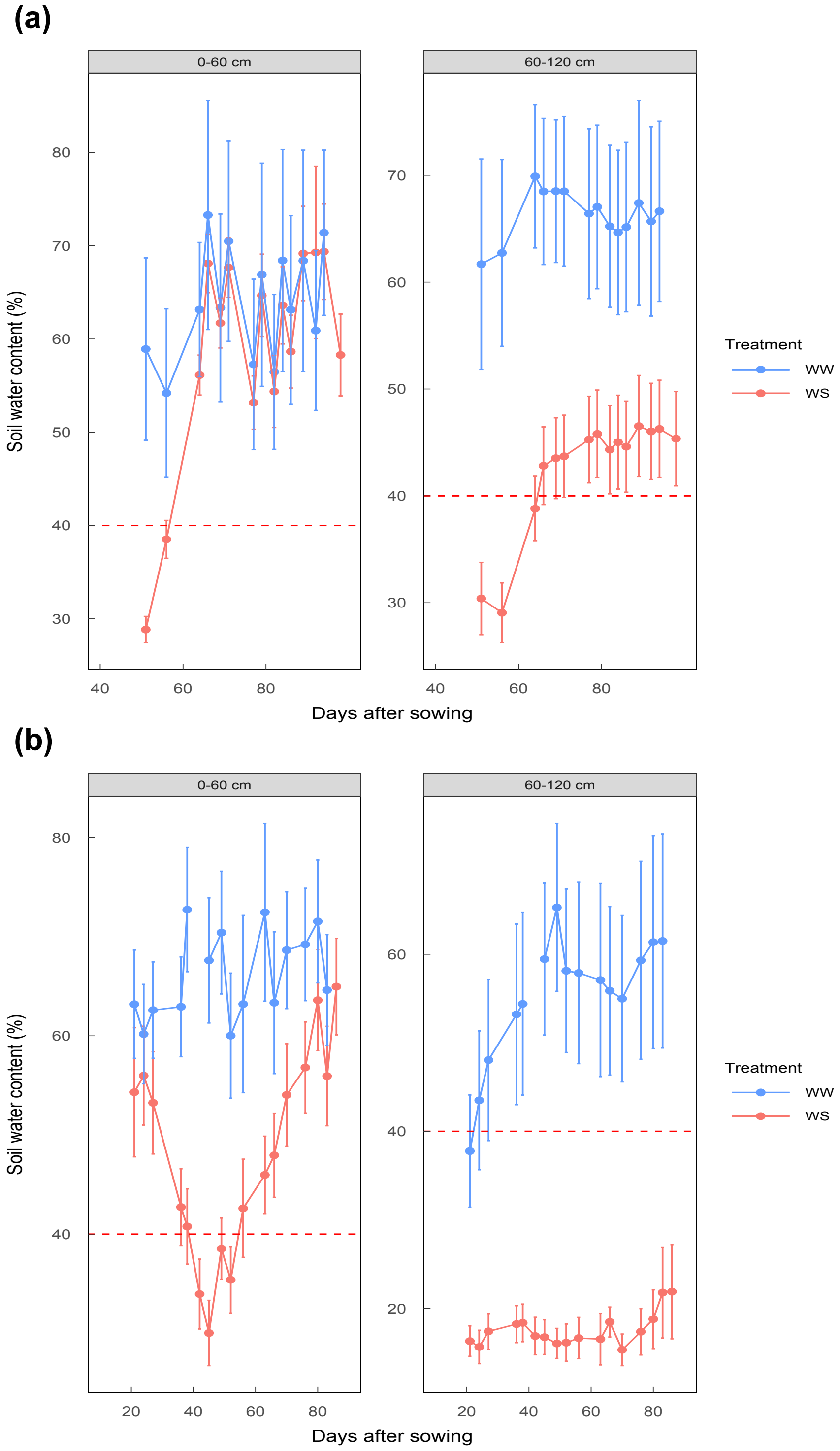


**Fig. S2** Volumetric soil water content in the 2021 (a, b) and 2022 (c, d) field trials at two depth intervals (0-60 cm and 60-120 cm) measured using DIVINER probes. The dry down started at 21 days after sowing (DAS) and irrigation was resumed at 49 days after sowing in 2021 and 42 days after sowing in 2022. In 2021, soil water content measurements started at 49 days after sowing. The red dotted line is considered as the onset of water stress. WW: Irrigated; WS: Drought stress.

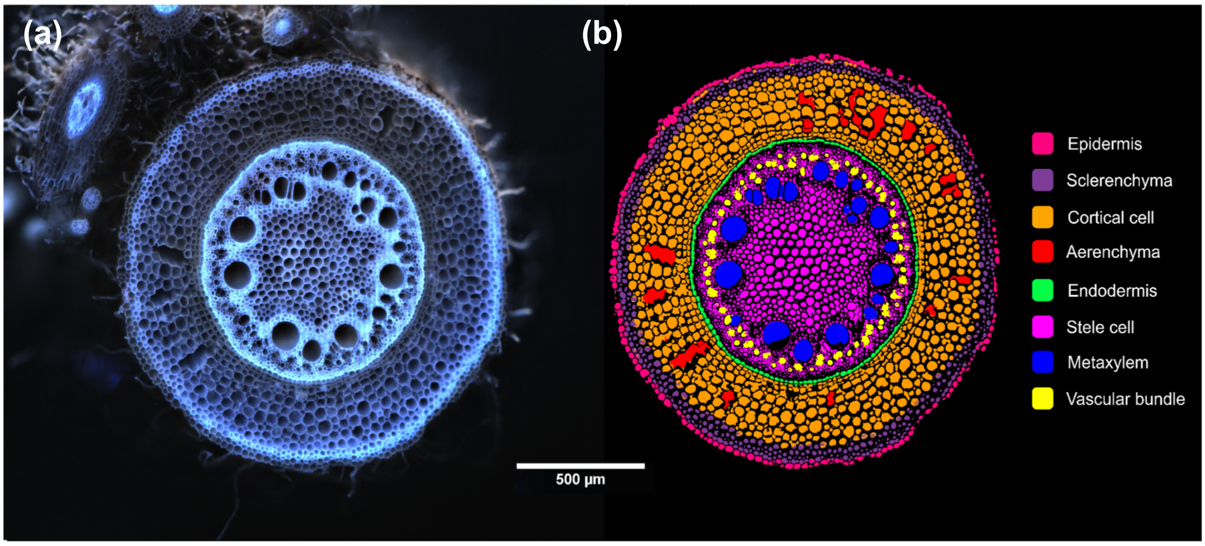


**Fig. S3** An example cross-sectional image of a crown root from node four obtained through laser ablation tomography (a) and tissue annotation using the pearl millet version of RootScan (b).

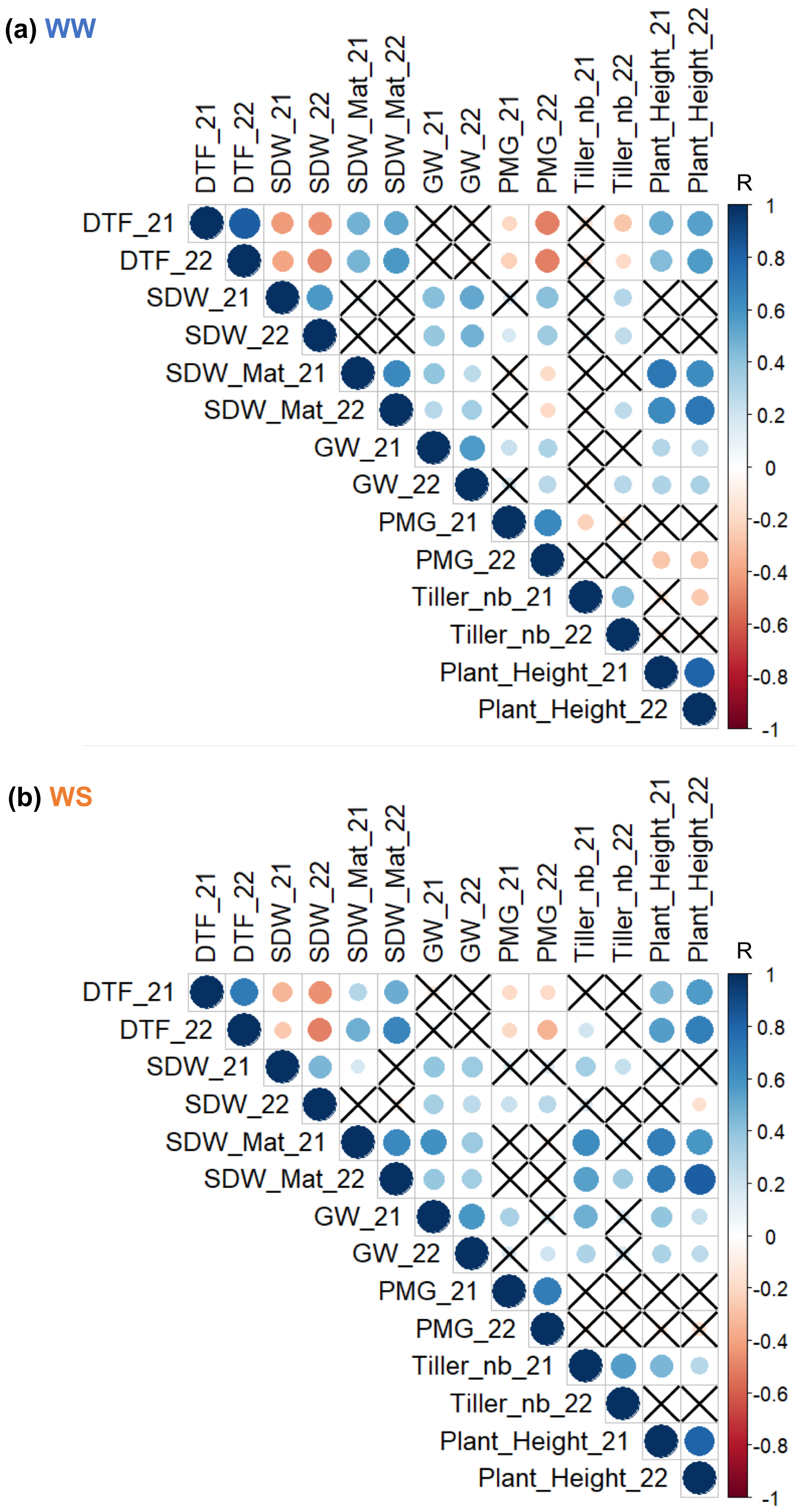


**Fig. S4** Correlation between shoot morphological and agronomical traits measured in the field across both years (2021 and 2022) in the irrigated (a, WW) and drought stress (b, WS) treatments. The Pearson correlation coefficient (R) is indicated for each pair of traits and black crosses indicate non-significant correlation (*p*-value > 0.05). DTF: Date to flowering; SDW_21: Shoot biomass at 49 days after sowing measured in the 2021 field experiment; SDW_22: Shoot biomass at 42 days after sowing measured in the 2022 field experiment; SDW_Mat: Shoot biomass at maturity; GW: Grain weight; PMG: 1000-grain weight; Tiller_nb: Tiller number; Plant_Height: Plant height.

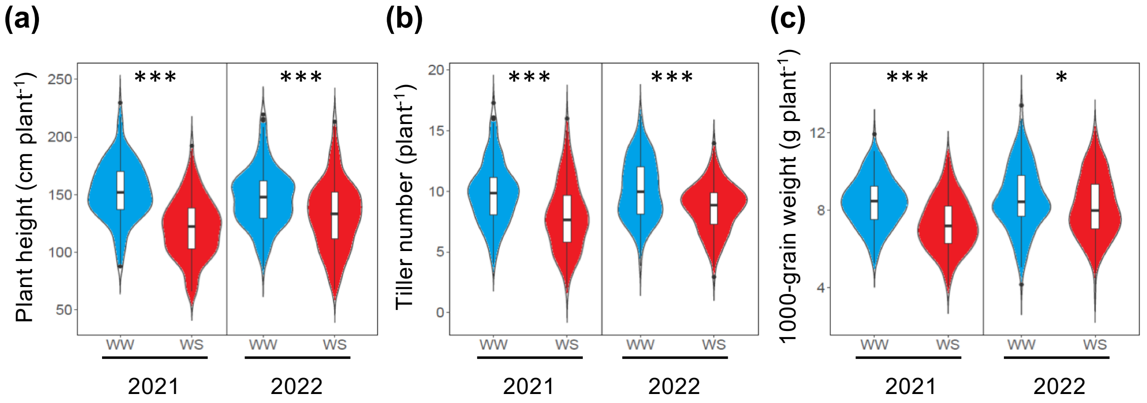


**Fig. S5** Stress impact on plant height (a), tiller number (b) and 1000-grain weight (c) measured in the field across both treatments (WW: Irrigated; WS: Drought stress) and years (2021 and 2022). Violin plots were produced using best linear unbiased estimates from all genotypes present within one treatment and year. * *p*-value < 0.05 and *** *p*-value < 0.001 indicate significant differences between treatments according to a Wilcoxon test.

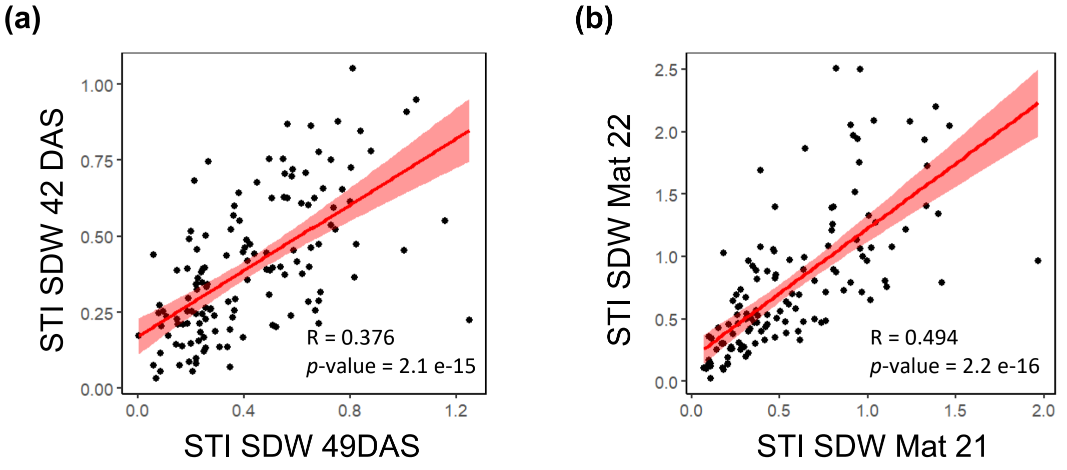


**Fig. S6** Covariation between stress tolerance index for shoot biomass measured in both years (2021 and 2022). (a) Stress tolerance index for shoot biomass measured at 49 days after sowing in 2021 (STI SDW 49 DAS) and at 42 days after sowing in 2022 (STI SDW 42 DAS). b) Stress tolerance index for shoot biomass measured at maturity (STI SDW Mat) in 2021 and 2022. The Pearson correlation coefficient (R) and *p*-value of the correlation test are indicated.

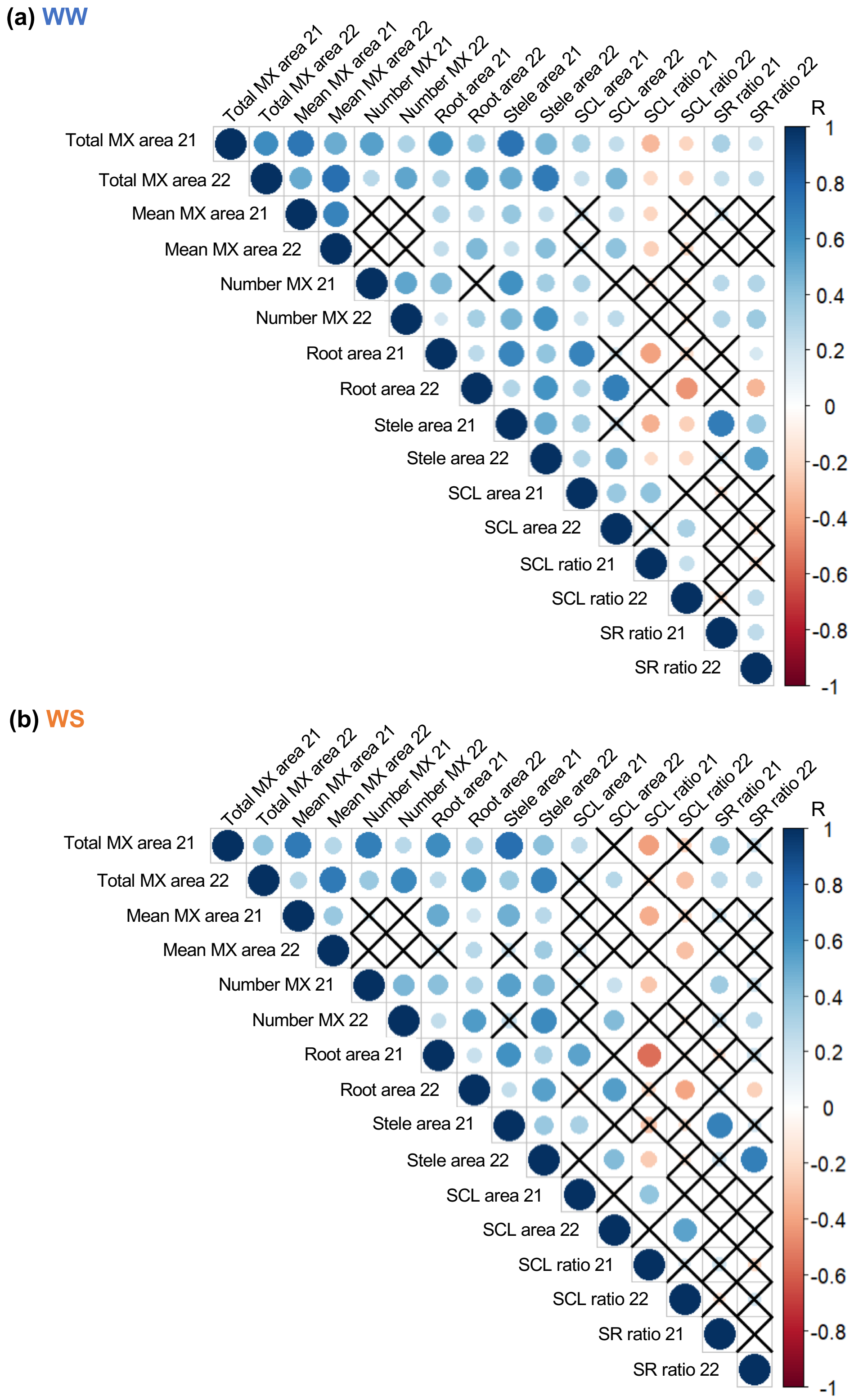


**Fig. S7** Correlation between root anatomical traits measured in the field across both years (2021 and 2022) in the irrigated (a, WW) and drought stress (b, WS) treatments. The Pearson correlation coefficient (R) is indicated for each pair of traits and black crosses indicate non-significant correlation ($p$-value > 0.05). MX: Metaxylem; Root area: Root cross section area; SCL: Sclerenchyma; SCL ratio: Ratio of sclerenchyma area to root cross section area; SR ratio: Ratio of stele area to root cross section area.

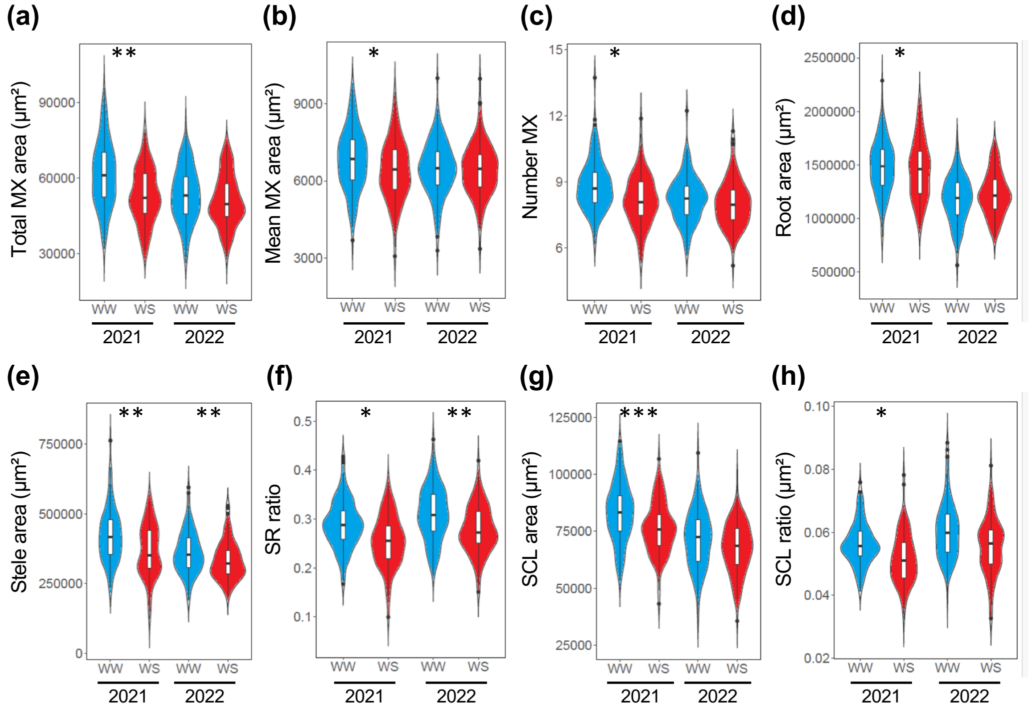


**Fig. S8** Stress impact on total metaxylem vessel area (a), mean area of metaxylem (b), number of metaxylem (c), root cross section area (d), stele area (e), ratio of stele area to root cross section area (SR ratio; f), sclerenchyma area (g) and ratio of sclerenchyma to root cross section area (SCL ratio; h). Root anatomical traits were measured in the field across both treatments (WW: irrigated; WS: Drought stress) and years (2021 and 2022). Violin plots were produced using best linear unbiased estimates from all genotypes present within one treatment and year. * $p$-value < 0.05, ** $p$-value < 0.01 and *** $p$-value < 0.001 indicate significant differences between treatments according to a Wilcoxon test. MX: Metaxylem; SCL: Sclerenchyma.

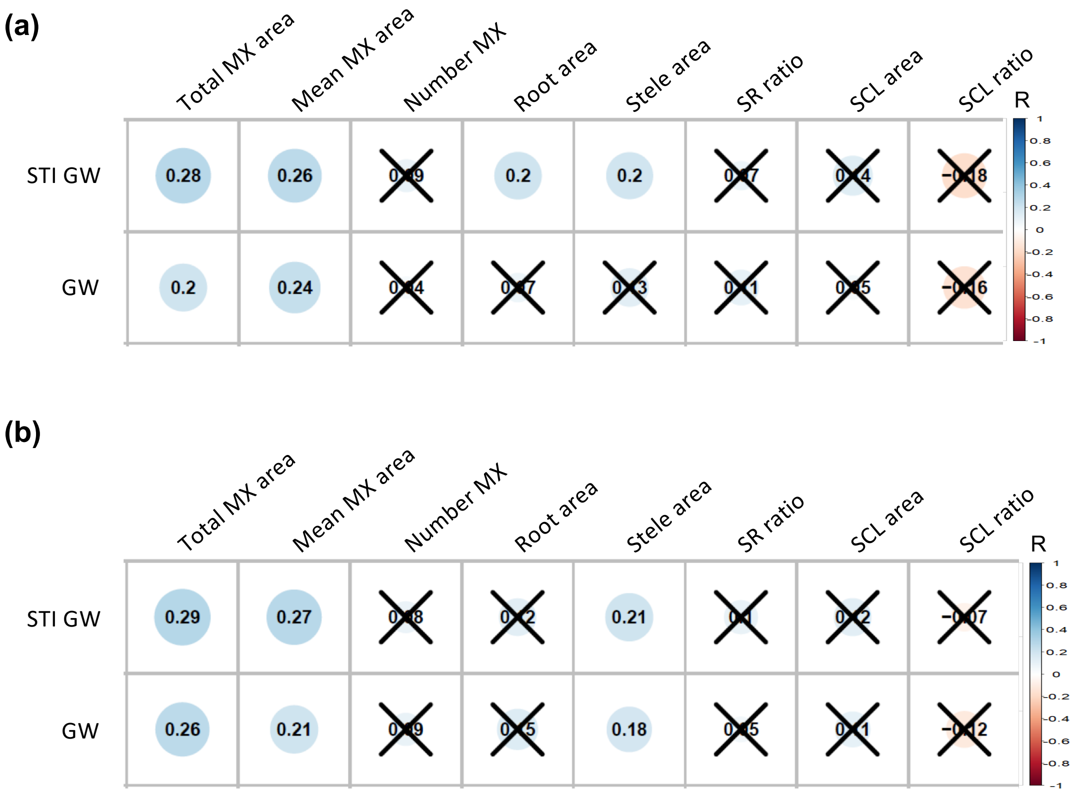


**Fig. S9** Correlation between root anatomical traits, grain weight (GW), and stress tolerance index for grain weight (STI GW) measured under irrigated treatment in the 2021 (a) and 2022 (b) field experiments. The Pearson correlation coefficient (R) is indicated for each pair of traits and black crosses indicate non-significant correlation (*p*-value > 0.05). MX: Metaxylem; SCL: Sclerenchyma; Root area: Root cross section area; SCL ratio: Ratio of sclerenchyma area to root cross section area; SR ratio: ratio of stele area to root cross section area.

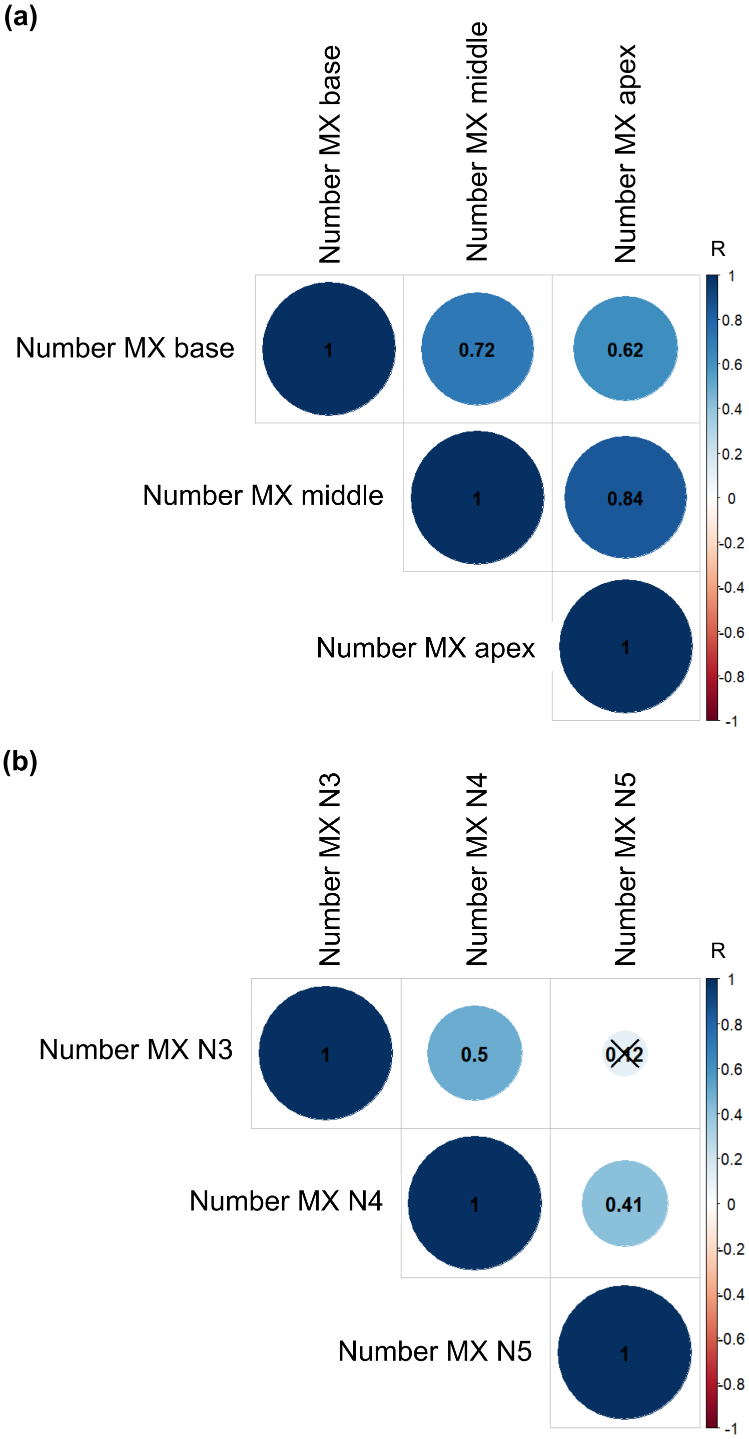


**Fig. S10** Correlation between the number of metaxylem vessels along the crown root of node four (a) and across different nodes (b). The Pearson correlation coefficient (R) is indicated for each pair of traits and black crosses indicate non-significant correlation ($p$-value > 0.05). MX: Metaxylem; N3: crown root from node 3; N4: crown root from node 4; N5: crown root from node 5.

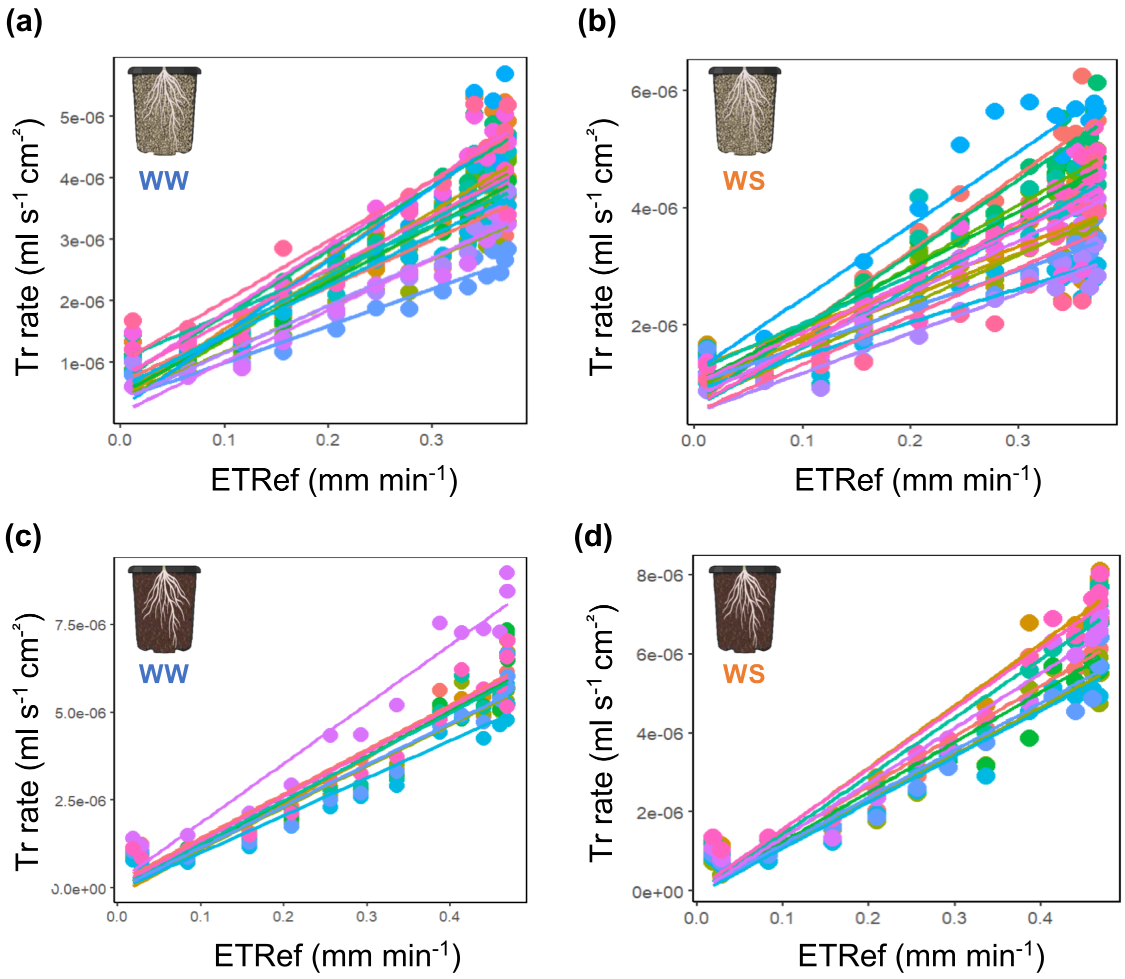


**Fig. S11** Transpiration response to the evaporative demand in pearl millet genotypes grown in the greenhouse under irrigated (WW) and drought stress (WS) treatments. (a, b) Plants were grown in sandy soil. (c, d) Plants were grown in peat soil. Tr rate: Transpiration rate; ETRef: Reference evapotranspiration measured in 30 minutes intervals.

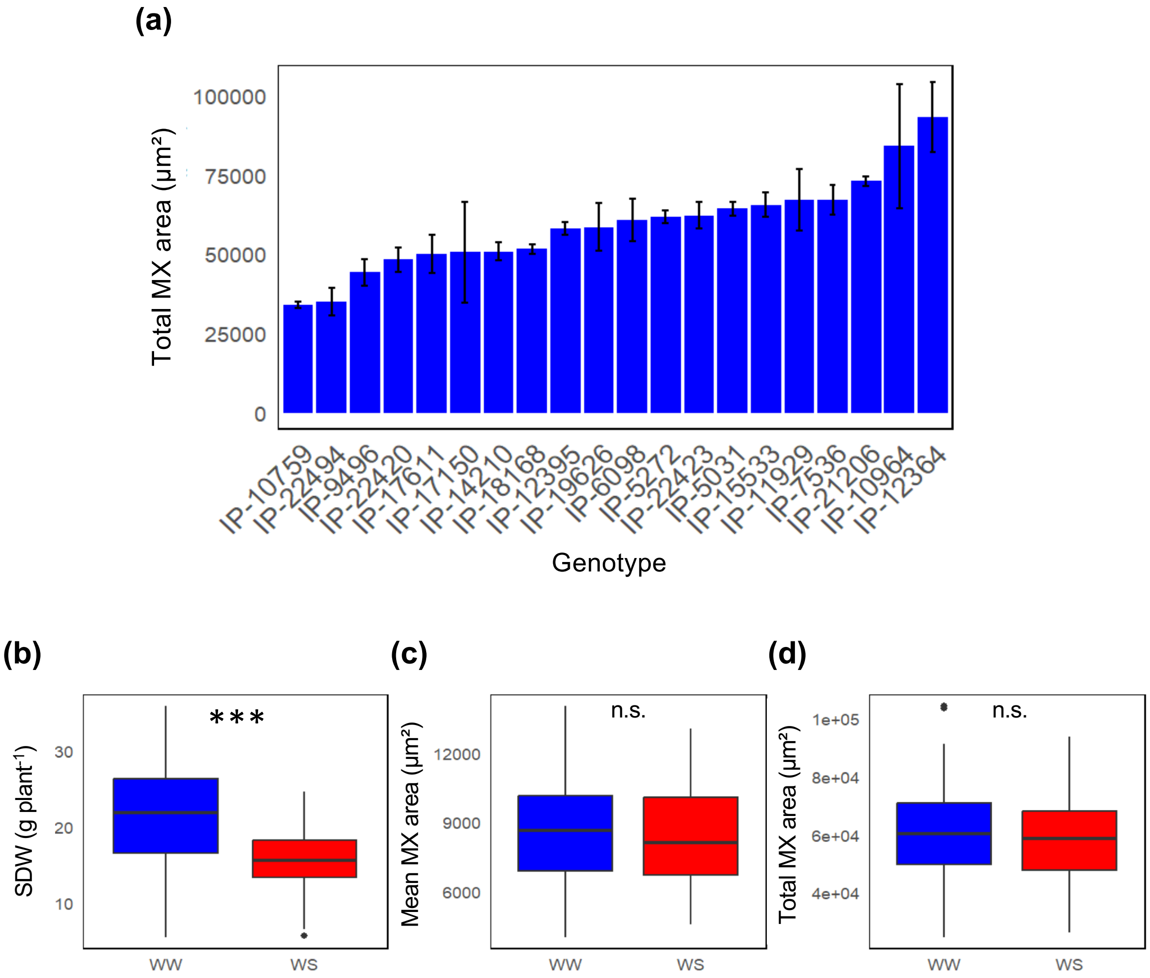


**Fig. S12** Stress impact on shoot biomass and metaxylem-related traits in pearl millet genotypes contrasting for total metaxylem area. Plants were grown in the greenhouse in sandy soil. (a) Total metaxylem area measured in the irrigated treatment. (b, c, d) Boxplots were produced using corrected averaged values from all genotypes present within one treatment. WW: Irrigated; WS: Drought stress; SDW: Shoot biomass; MX: Metaxylem; *** $p$-value < 0.001 indicates significant differences between treatments according to a Wilcoxon test. n.s.: Not significant.

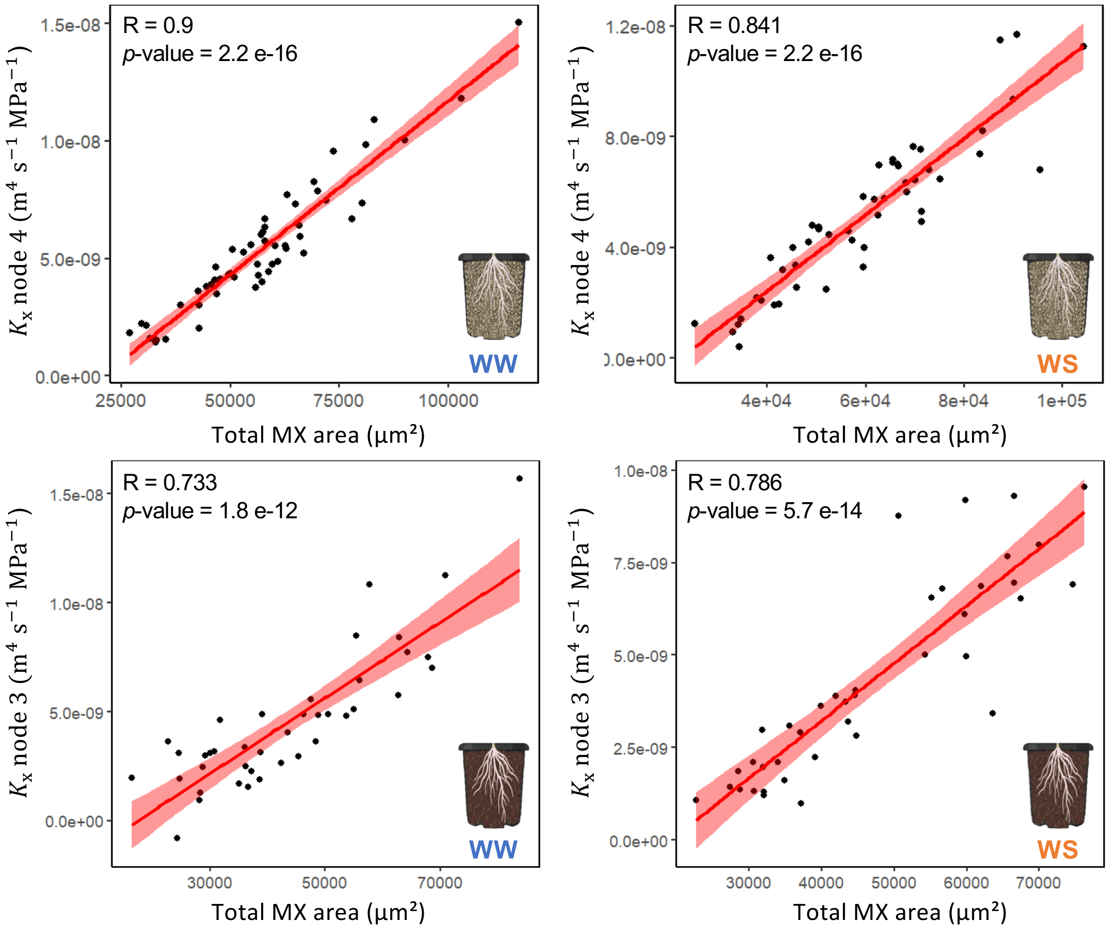


**Fig. S13** Covariation between total area of metaxylem and axial root hydraulic conductance. (a, b) Plants were grown in sandy soil in the greenhouse. Total metaxylem area and axial root hydraulic conductance were measured on crown roots from node 4 under irrigated (WW; a) and drought stress treatments (WS; b). (c, d) Plants were grown in peat soil in the greenhouse. Total metaxylem area and axial root hydraulic conductance were measured on crown roots from node 3 under irrigated (WW; c) and drought stress treatments (WS; c). MX: Metaxylem.

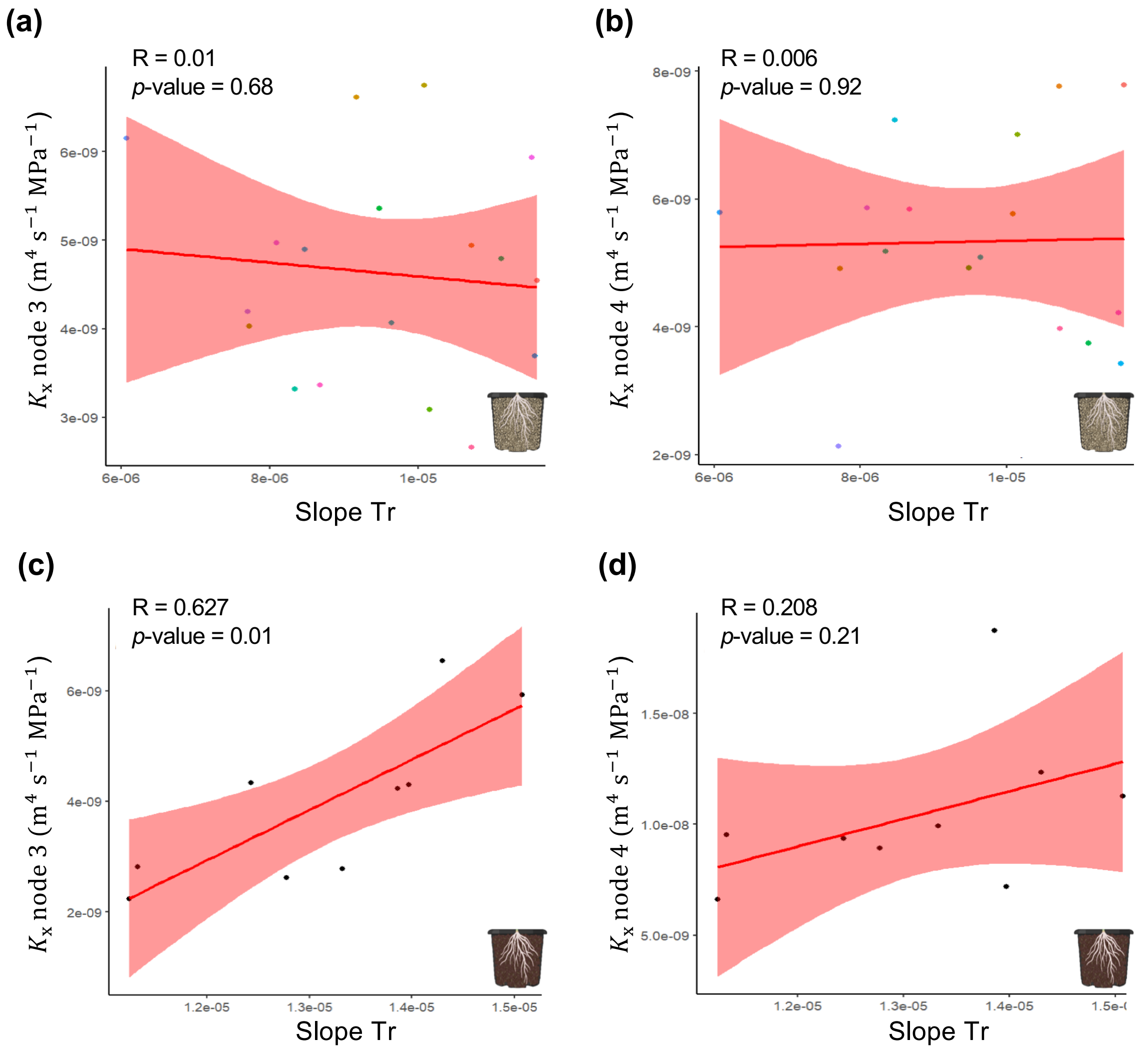


**Fig. S14** Covariation between axial root hydraulic conductance ($K_x$) measured in crown roots from node three and four, and the transpiration response to the evaporative demand (Slope Tr) in pearl millet genotypes contrasting for total metaxylem area grown under irrigated treatment in the greenhouse. (a, b) Plants were grown in sandy soil. (c, d) Plants were grown in peat soil. The Pearson correlation coefficient (R) and *p*-value of the correlation test are indicated.

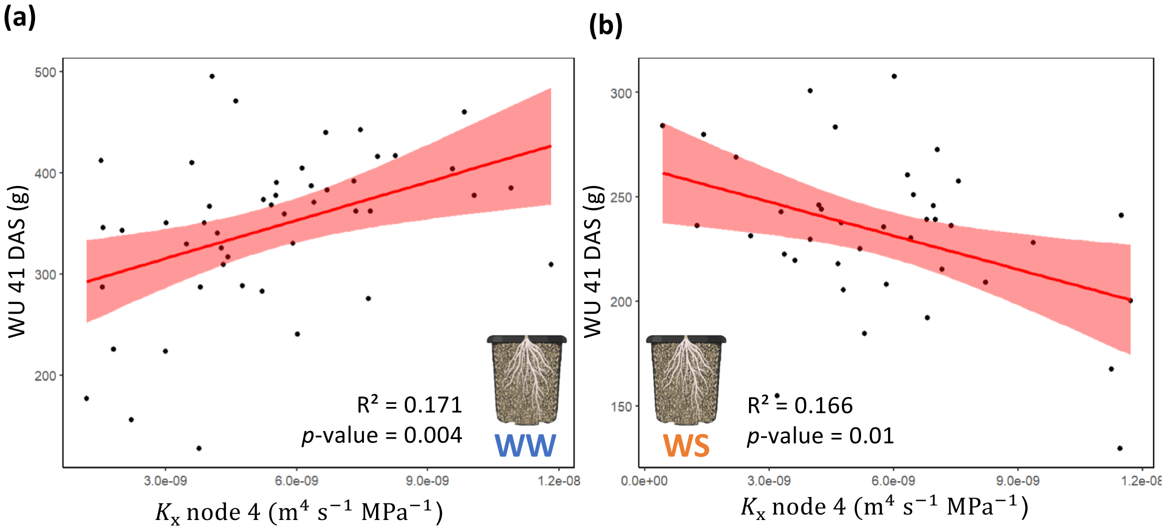


**Fig. S15** Water use in pearl millet genotypes contrasting for total metaxylem area grown under irrigated (WW) and drought stress (WS) treatments in sandy soil in the greenhouse. (a, b) Covariation between the cumulated water use at 41 days after sowing and axial root hydraulic conductance ($K_x$) measured on crown root from node four in all genotypes under irrigated (a) and drought stress (b) treatments. The Pearson correlation coefficient (R) and *p*-value of the correlation test are indicated. WW: Irrigated; WS: Drought stress.

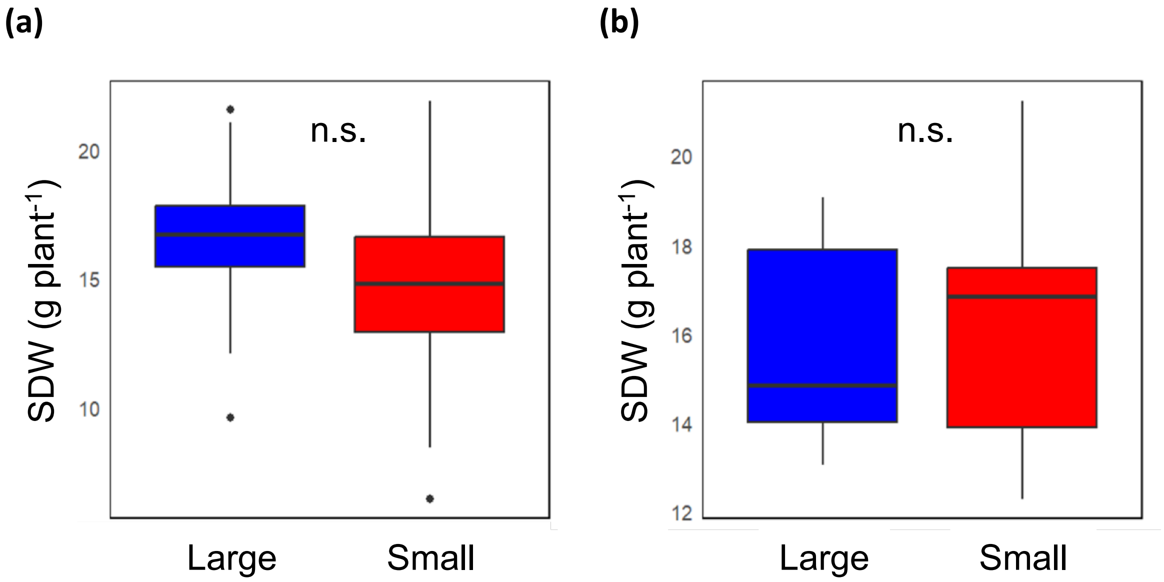


**Fig. S16** Shoot biomass in two groups of genotypes contrasting for axial root hydraulic conductance ($K_x$; large versus small) measured under the drought stress treatment. (a) Plants were grown in sandy soil. (b) Plants were grown in peat soil. n.s.: Non significant differences were observed between the two groups according to a Wilcoxon test.

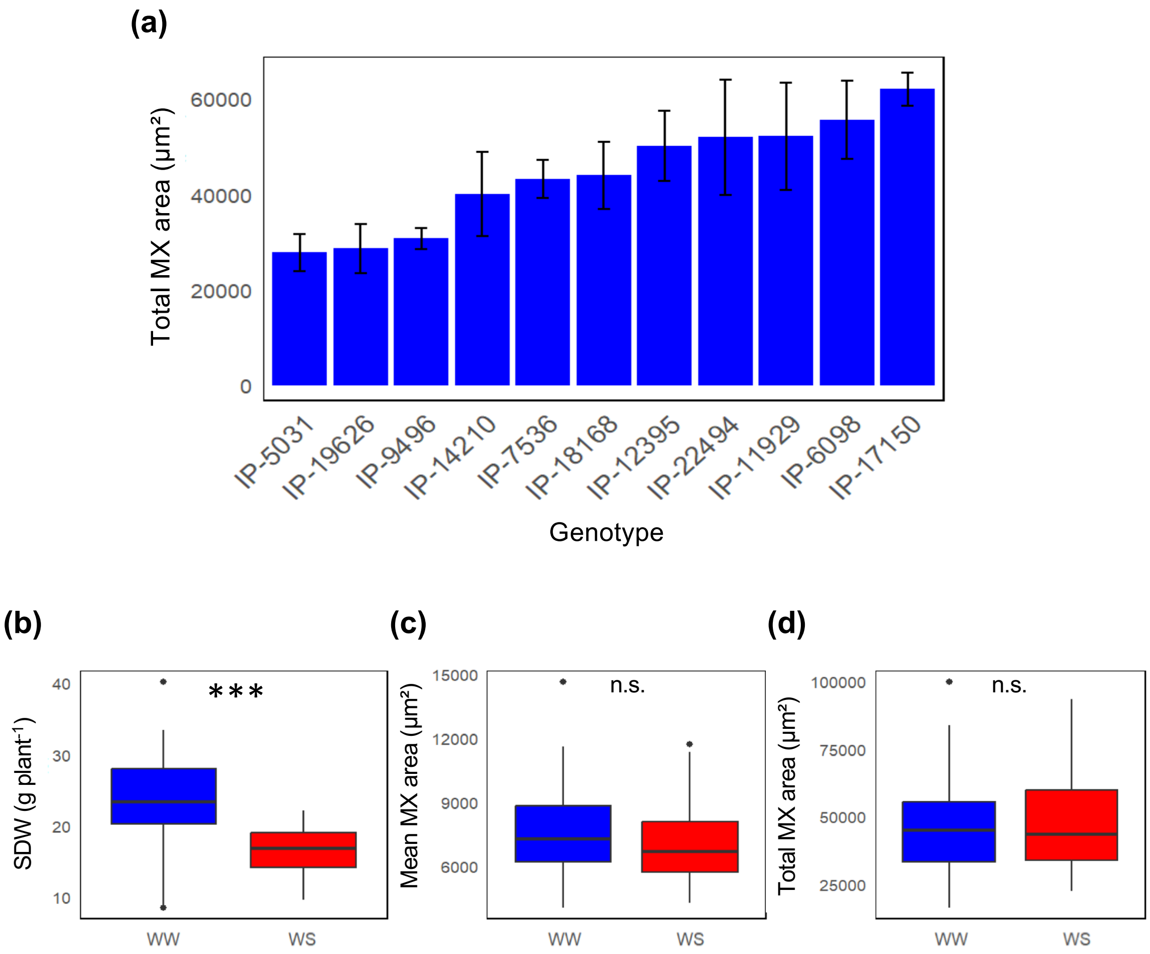


**Fig. S17** Stress impact on shoot biomass and metaxylem-related traits in pearl millet genotypes contrasting for total metaxylem area. Plants were grown in the greenhouse in peat soil. (a) Total metaxylem area measured in the irrigated treatment. (b, c, d) Boxplots were produced using corrected averaged values from all genotypes present within one treatment. WW: Irrigated; WS: Drought stress; SDW: Shoot biomass; MX: Metaxylem; *** *p*-value < 0.001 indicates significant differences between treatments according to a Wilcoxon test. n.s.: not significant.

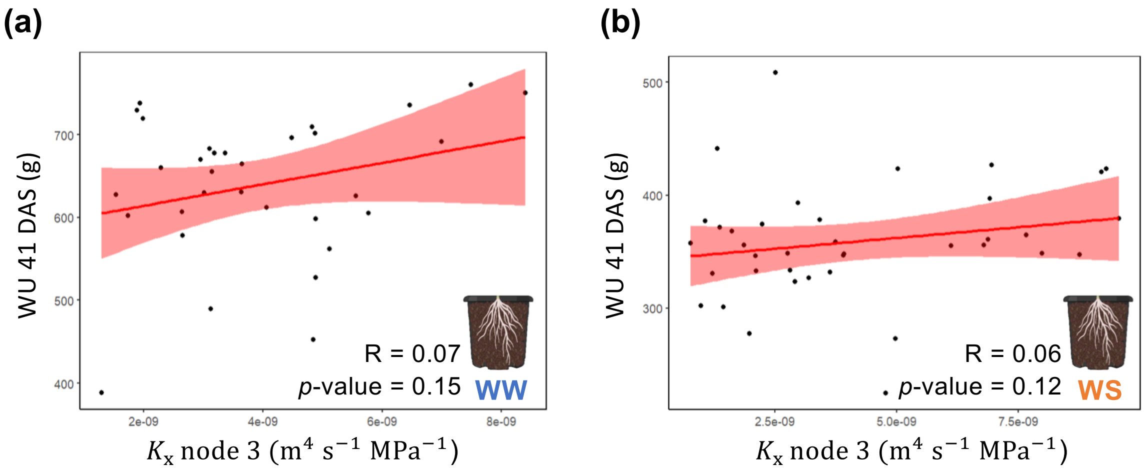


**Fig. S18** Water use in pearl millet genotypes contrasting for total metaxylem area grown under irrigated (WW) and drought stress (WS) treatments in peat soil in the greenhouse. (a, b) Covariation between the cumulated water use at 41 days after sowing and axial root hydraulic conductance ($K_x$) measured on crown root from node four in all genotypes under irrigated (a) and drought stress (b) treatments. The Pearson correlation coefficient (R) and *p*-value of the correlation test are indicated. WW: Irrigated; WS: Drought stress.

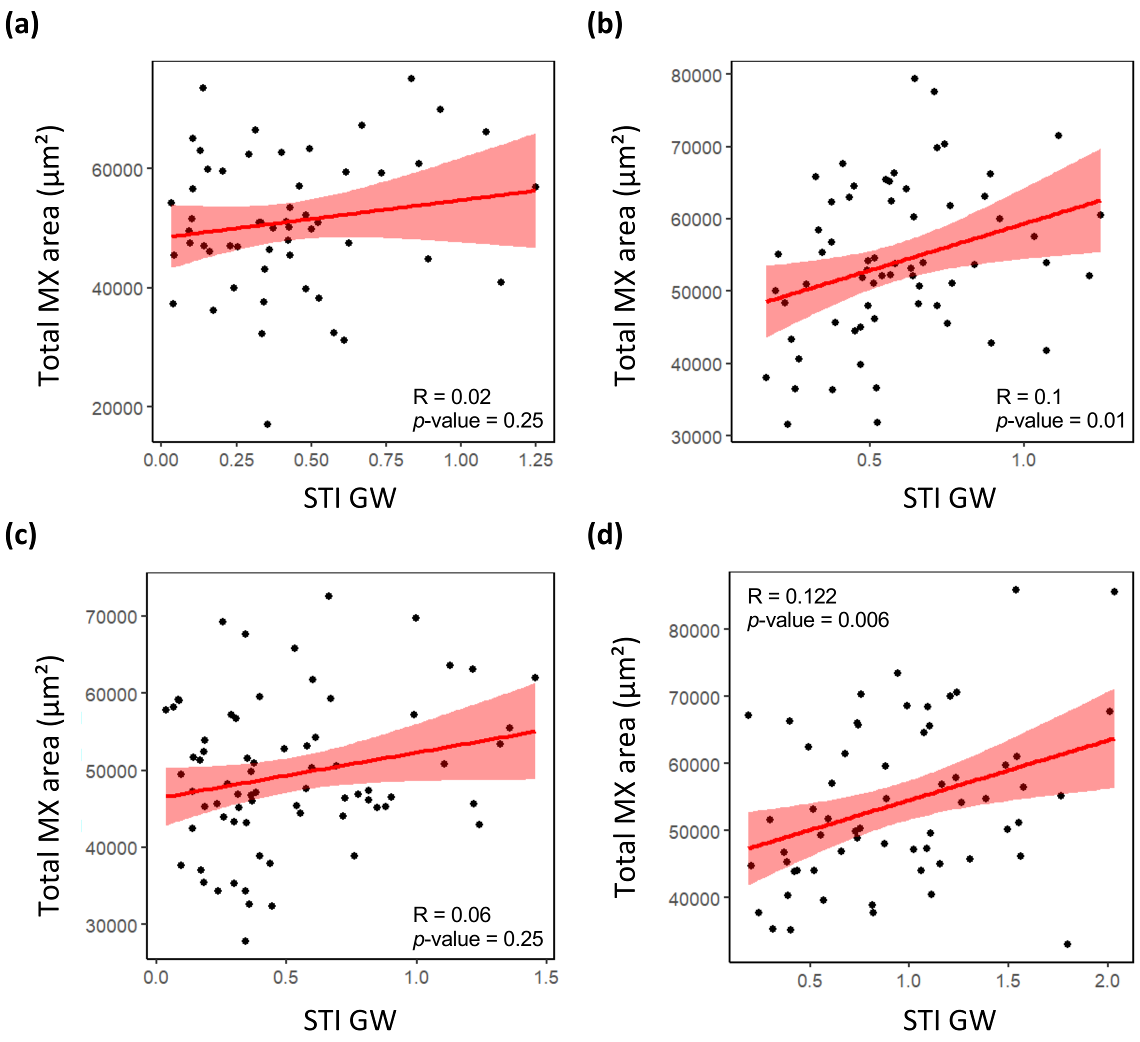


**Fig. S19** Covariation between total metaxylem area and stress tolerance index for grain weight. Covariations were performed within two shoot biomass groups measured at 49 days after sowing (2021; a, b) or 42 days after sowing (2022; c, d) based on a clustering analysis. (a, c) Covariations within the low shoot biomass group. (b, d) Covariations within the high shoot biomass group. MX: Metaxylem; STI GW: Stress tolerance index for grain weight.

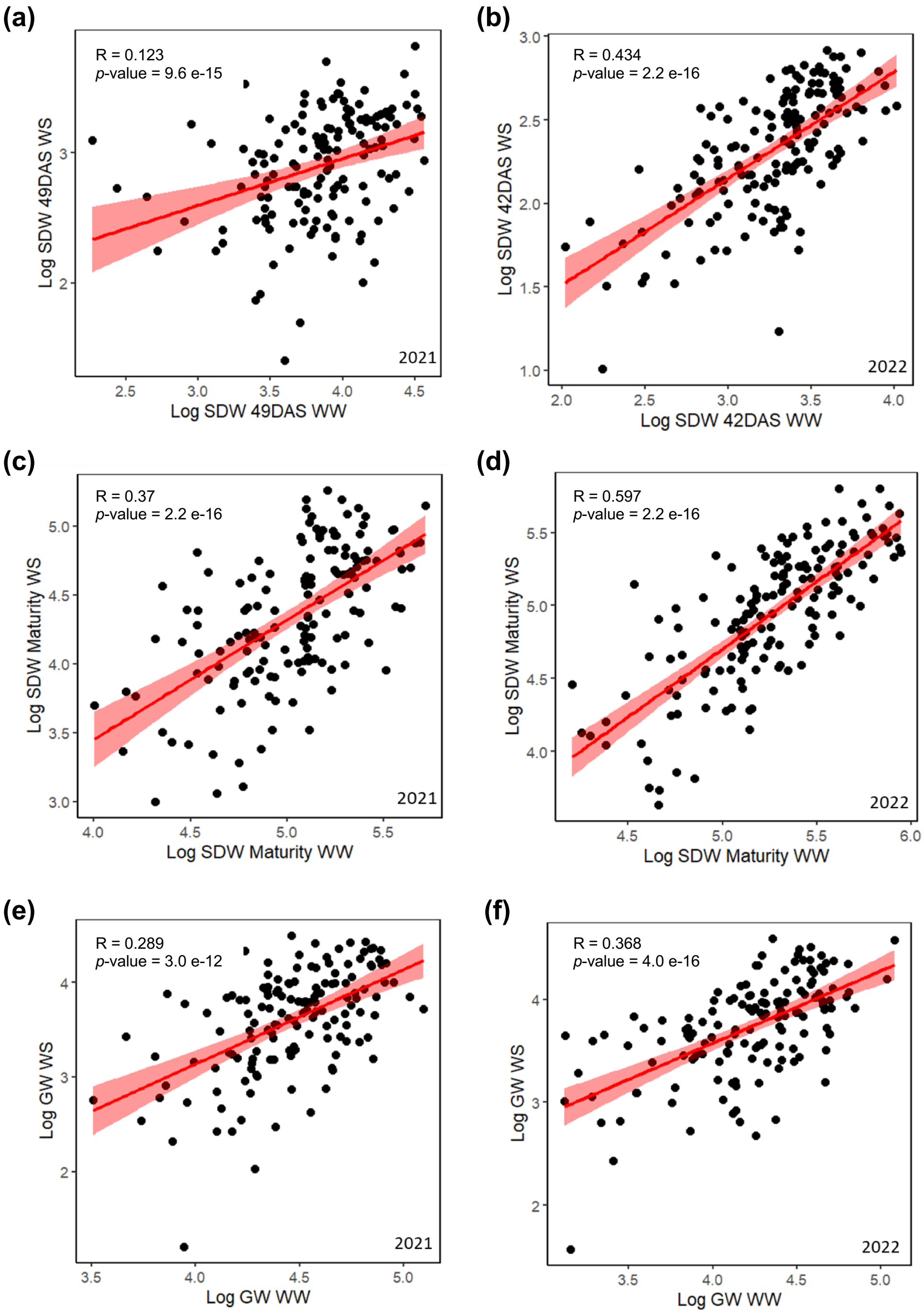


**Fig. S20** Covariation of shoot biomass and grain yield between both treatments within years. (a, b) Covariation of shoot biomass measured at the end of the drought stress in 2021 (49 days after sowing; a) and in 2022 (42 days after sowing; b) between the irrigated (WW) and drought stress (WS) treatments. (c, d) Covariation of shoot biomass measured at maturity in 2021 (c) and in 2022 (d) between the irrigated (WW) and drought stress (WS) treatments. (e, f) Covariation of grain weight in 2021 (e) and in 2022 (f) between the irrigated (WW) and drought stress (WS) treatments. GW: Grain weight; SDW: Shoot biomass. Values were expressed on a logarithmic scale. The Pearson correlation coefficient (R) and *p*-value of the correlation test are indicated.
